# The Network Basis of Pattern Formation: A Topological Atlas of Multifunctional Turing Networks

**DOI:** 10.1101/2025.01.27.634997

**Authors:** Laura Regueira López de Garayo, Luciano Marcon

**Affiliations:** Centro Andaluz de Biología del Desarrollo (CABD) Universidad Pablo de Olavide-CSIC-JA Carretera de Utrera Km.1 41013, Seville, Spain

**Keywords:** Turing, Self-Organization, Network Topology, Pattern-formation, Noise

## Abstract

Understanding how genetic networks can drive different self-organizing spatial behaviors remains a significant challenge. Here, we use an automated algebraic method to systematically screen for Turing networks capable of generating diverse spatial patterns from noise, including periodic static waves, traveling waves and noise-amplifying patterns. We organize these networks into a topological atlas—a higher-level graph where nodes represent Turing networks linked together when they differ by only one regulatory interaction. In this atlas, Turing networks are arranged into distinct clusters showing a remarkable correspondence between network topology and self-organizing behaviors. Using an analytical approach, we identify the specific regulatory feedbacks that characterize each behavior. Moreover, we discover that different clusters are interconnected by multifunctional networks that can transition between behaviors upon feedback modulation. Among these networks, we find a new class of multiphase Turing networks capable of altering the phase of periodic wave patterns depending on the parameters, and networks that can transition between static and oscillatory Turing behaviors. The atlas further highlights the crucial role of feedback on immobile nodes in regulating pattern formation speed and precision by canalizing system noise. Overall, our study provides a novel framework to study the evolution and development of multicellular self-organization through changes in network topology and feedback modulation. This offers insights into how genetic regulatory networks can be tuned to drive pattern formation in developmental biology and in stem cell systems like embryoids and organoids.

**Significance Statement:** By employing an automated algebraic method, Regueria and Marcon construct a topological atlas that categorizes Turing networks based on their ability to produce periodic patterns, traveling waves, or noise amplifying patterns. The atlas identifies distinct topological clusters linked by multi-functional networks that can transition between behaviors through feedback modulation. Key findings highlight how modulation of regulatory cycle strength in time or space can promote transition between static and oscillatory periodic pattern. The study also reveals the importance of feedback on immobile nodes in managing noise and influencing pattern formation. Overall the topological atlas offers a new framework for examining the evolution and development of multicellular self-organization.

---

**T**uring’s reaction-diffusion model explains how uniform systems can break symmetry to generate spatial patterns. Initially proposed to explain early symmetry-breaking and morphogenesis of the embryo (1), the model was overlooked in favor of hierarchical systems such as positional information (2). Recently, interest in Turing’s theory has been renewed in developmental biology (3) and for studying the self-organization of embryoids and organoids (4). Despite this resurgence, the precise genetic interactions driving different type of Turing patterns in multicellular systems remain largely unknown.

Turing proposed that cells could self-organize by exchanging substances called morphogens, which diffused between cells like hormones and interacted according to standard chemical reactions. Depending on reaction terms, morphogen systems could form six types of spatial waves categorized as stationary or oscillatory with extremely long, extremely short, or finite wavelengths (1). Patterning dynamics resembling static or oscillatory Turing patterns have been observed in various biological systems like skin appendages and limb development (5, 6). Although the key genes involved in these patterning events have been identified, understanding the interactions that drive Turing behaviors requires analysis through regulatory principles rather than chemical stoichiometry, as originally proposed by Turing.

Theoretical network screenings have systematically analyzed gene regulatory networks, identifying simple regulatory principles that promote oscillations (7) and adaptive responses (8, 9). A comprehensive screening (10) identified three-node networks capable of forming a peak of gene expression in response to a gradient. This study organized the networks into an atlas constructed as a higher-level graph of networks, where each node represented a gene regulatory topology, and edges connected topologies differing by only one interaction (10). This was a convenient approach to organizing networks in topological space, clustering similar topologies together, and identifying six minimal regulatory motifs linked to variations in underlying feed-forward logic (11). A subsequent study (12) expanded this atlas to include other patterning behaviors, finding that topological regions of different mechanisms were connected via multifunctional networks that could switch behaviors depending on parameters.

Theoretical network screenings, however, have not been traditionally applied to Turing systems. Instead, Turing systems were often studied using minimal two-species models (3, 13) due to the complexity of deriving patterning conditions for larger networks (1, 14–20). An exception to this were pioneering studies that performed a random numerical screening for all gene regulatory networks capable of spatial pattern formation (21, 22). More recently, an automated computer algebra approach was used to analyze larger Turing networks systematically, identifying minimal three- and four-node networks with one and two immobile nodes respectively (23). This was achieved by deriving analytical conditions to determine the signs of the solutions of the characteristic polynomial obtained by linear stability analysis (1, 24). These solutions relate the eigenvalue to the potential spatial patterns (wavenumbers), forming what is known as the dispersion relation. This analysis confirmed that the network’s structure depicted by the Jacobian matrix, which illustrates how substances interact around a stable state according to the linear stability analysis, effectively predicts the pattern-forming capabilities of reaction-diffusion systems (23, 25).

A subsequent study used parameter sampling for linear stability analysis of 2- and 3-node networks (26), performing numerical screening to identify parameters yielding a positive eigenvalue instead of solving analytically the linear stability analysis. While offering limited parameter coverage, this method allowed scaling up to non-minimal 3-node networks with more than six interactions. It also considered cases where all three nodes were diffusible. The model proposed that all 3-node networks reduced to two types of 2-node networks (AIJT and CAIJT), corresponding to minimal Jacobian signs for Turing patterns, referred to in (13) as Activator-Inhibitor and Substrate-Depletion systems. The study indicated greater robustness in AIJT networks, though this depended on the specific nonlinearities and the implementation of the core topologies. Indeed, in the simple linear case, these two topologies have the same parameter space size for Turing patterns (24).

Another parameter sampling approach represented each network according to the reaction terms in partial differential equations, distinguishing competitive and non-competitive interactions (27). This generated a broader network list but introduced ambiguity, as it did not clarify which terms dominate around the homogeneous steady state. For example, unlike in the Jacobian-based representation, negative linear terms were ignored in these network diagrams. This led to the proposal that five different 2-node networks can make Turing patterns, while traditional Jacobian-based methods identify only two (1, 23, 24, 26, 27). The study proposed that Turing networks are sensitive to parameter variations indicating low robustness. It also identified that core regulatory motifs such as positive feedback on diffusing nodes, diffusion-mediated negative feedback loops, and competitive interactions were prevalent in robust Turing networks (27).

While previous screenings attempted to relate Turing networks to identify regulatory principles (26, 27), they did not generated a fully connected topological atlas that mapped all networks, as achieved in (10, 12). Moreover, these studies primarily focused on Turing networks generating static periodic patterns, neglecting oscillatory patterns (23, 26, 27) and noise-amplifying networks. These networks, also known as Turing filters, were first described in (23, 25) and later identified in (27). Turing filters meet Turing conditions but have dispersion relations that exhibit asymptotic behaviour for large wavenumbers, as first described in (16). This results in the amplification of all spatial patterning modes present in the initial conditions, leading to noisy patterns from random initial conditions (23, 27) or periodic spatial patterns from localized initial conditions (25, 28). A crucial requirement for this behaviour is the absence of a maximum eigenvalue in the dispersion relation. When the dispersion relation has a maximum eigenvalue peak, the presence of a lower positive asymptotic behaviour does not interfere with the Turing network’s ability to form patterns from noise, as initially described in (23, 25, 29) and later confirmed in (27).

In this study, we extend our previous analytical screening approach (23) to identify networks that generate both static and oscillatory Turing patterns, and also consider noise-amplifying networks. We organized these networks into a fully connected atlas, allowing transitions between Turing networks and behaviours by systematically adding or removing single interactions. By examining transitions within the atlas and exploiting the formulas from our analytical approach, we identify multiphase and multifunctional networks that switch between pattern phase relations and behaviours based on regulatory feedback changes. Additionally, we find that feedbacks on immobile nodes control noise canalization, which is crucial for pattern timing and precision.

Overall, we show that the atlas helps understand how regulatory feedback modulation in Turing networks promotes transitions between self-organizing patterning behaviors during evolution or development. This approach contributes to translate Turing’s chemical basis of morphogensis into a framework based on regulatory network feedback, which is more suitable for studying multicellular pattern formation driven by genetic networks.

## Results

We performed a comprehensive analysis to construct and understand a topological atlas of 3-node Turing networks with one immobile node. This analysis focuses on identifying networks that can generate static and oscillatory diffusion-driven patterns, as well as noise-amplifying networks. This was done in a completely algebraic manner without relying on parameter sampling and numerical simulations, but rather by deriving the conditions for the existence of positive or negative real roots λ of the characteristic polynomial *P* (λ) = λ^3^ + *a*_1_(*q*)λ^2^ + *a*_2_(*q*)λ + *a*_3_(*q*) in each network, where the coefficient (*a*_1_, *a*_2_, *a*_3_) contain symbolic parameters for the rates or network cycle weights, and diffusion coefficients. More details are provided in the Material and Methods.

By examining transitions between neighbouring networks, we identify the regulatory mechanisms driving different self-organizing behaviours analytically. Our results reveals that networks with distinct behaviors are linked by Multifunctional networks capable of transitioning between behaviors depending on feedback modulation. The following sections provide details on the atlas construction and an analysis of three paths along the atlas, revealing different properties of Multifunctional networks.

### Construction of the Topological Atlas

To construct the atlas, we began considering only minimal 3-node Turing networks having six regulatory interactions. We observed that changing the sign of a single interaction in these minimal networks (e.g., *k*_7_ in Figure 1A) disrupts Turing behavior and it is necessary to change the sign of two interactions simultaneously (e.g., *k*_2_ and *k*_3_) to maintain it, see SI Appendix. Since reconciling two simultaneous changes with the progressive changes that may occur during evolution is difficult, we decided to construct the atlas alternatively by including non-minimal (extended) networks with seven regulatory interactions. This approach allowed us to add or remove single interactions at the time (e.g., *k*_6_ in Figure 1B top) while preserving Turing behavior. This strategy resulted in a fully connected atlas where nodes represent Turing networks that are directly connected when differing by one interaction. Minimal networks with six edges are represented as square nodes, while those with seven edges are shown as circular nodes (Figure 1B), with paths in the atlas corresponding to sequences of alternating minimal and extended networks.

**Fig. 1.**
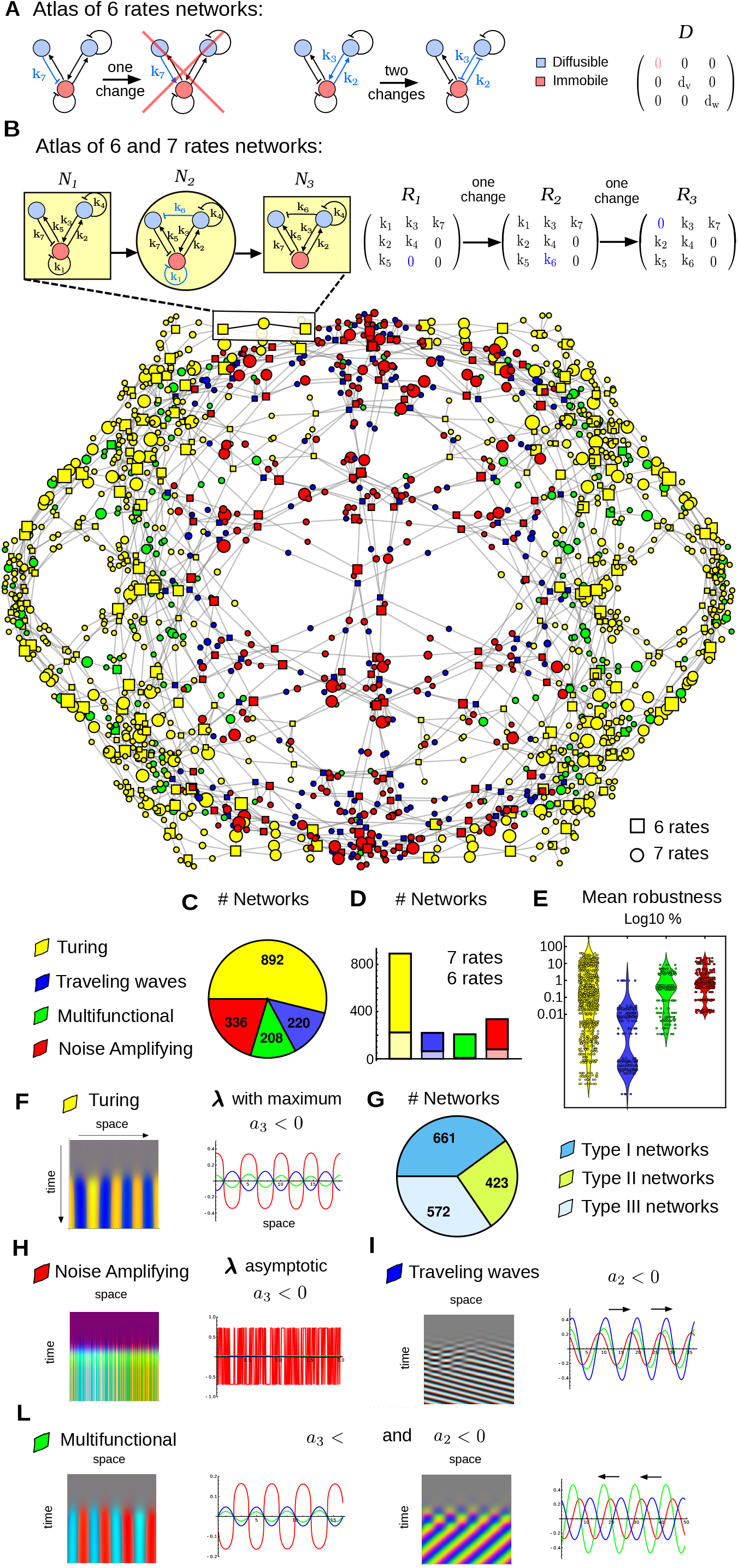
Topological atlas of 3-node Turing networks. A) A minimal 3-node Turing network with 6 interactions. Changing the sign of any interaction (e.g. k7) disrupts Turing behavior (red cross); two interactions must be changed simultaneously (e.g. k2, k3) to maintain Turing patterning. B) Adding a new single interaction (e.g. k6) can maintain Turing behavior. This strategy can be used to form a topological atlas, where nodes (e.g., *N*_1_, *N*_2_, *N*_3_) represent Turing networks. Node are connected when network differ by one interaction. Square nodes correspond to 6 edges networks (e.g., N1, N3), circular nodes with 7 edges (e.g., N2). Node size is proportional to the robustness of the network to parameter changes. Node color correspond to the of Turing behaviour exhibited by the network: Static Turing waves (yellow), Traveling waves (blue), Multifunctional: static or traveling waves depending on parameters (green) and Noise Amplification (red). C) Top: pie chart show the number of network for each behaviour. Number of 6 edges and 7 edges networks for each type of behaviour: 6 edges (light color) and 7 edges (dark color). Light green is very small because there are only 8 minimal networks out of 208 multifunctional networks. Mean network robustness to parameter space changes for each Turing behavior. Details of the calculation are provided in Materials and Methods. The logarithmic scale emphasizes the substantial differences in robustness across networks. Turing networks (yellow) are the most robust, with up to 40% of the parameter space capable of generating a diffusion-driven instability. These are followed by Noise (red), Multifunctional (green), and Traveling Wave networks, which exhibit at most 1% of the parameter space that promote a diffusion-driven instability. F) Simulation of a static Turing pattern network identified by a positive real λ with a maximum promoted by diffusion-driven instability when the coefficient *a*_3_ of the characteristic polynomial is negative. G) Number of Turing networks Type I (requires differential diffusivity), Type II (allow equal diffusivity) and Type III (any diffusivity). H) Simulation of a Noise Amplifying network identified by a positive real λ with an asymptotic behaviour promoted by diffusion-driven instability when *a*_3_ < 0. I) Simulation of a Traveling wave network identified by a complex positive λ promoted by diffusion-driven instability when *a*_1_ < 0 or *a*_2_ < 0. L) Simulations of a Multi-functional network that can form static Turing waves or Traveling waves depending on parameters.

Node size represents the robustness to parameter changes of the associated network, calculated as the portion of parameter space that gives rise to the self-organizing patterning behaviors. This was determined through a multiple integral over the defined parameter space for each network (see Material and Methods). Node colors represent the type of Turing behavior: static Turing waves (yellow), traveling waves (blue), multifunctional (green), and noise amplification (red).

Each network was categorized according to the sign of the characteristic polynomial’s coefficients *a*_1_, *a*_2_, and *a*_3_, which predict network behavior (see Materials and Method). If *a*_3_ < 0 and λ has no maximum, the network amplifies noise. If λ has a maximum, it corresponds to static Turing wave patterns. Having *a*_1_ < 0 or *a*_2_ < 0 is a sufficient (but not necessary) condition for networks to have a λ with a positive complex part, giving rise to traveling wavesm (see Material and Methods). Networks with both *a*_3_ < 0 and *a*_1_ < 0 or *a*_2_ < 0 are Multifunctional, capable of both static and oscillatory behavior depending on parameters.

Our analysis revealed that the majority of networks in the atlas generate static Turing patterns (yellow), followed by Noise-amplifying networks (red), waves (blue), and a minority of Multifunctional networks (green) (Figure 1C).

It also showed that extended networks are the majority in each class (Figure 1D). Multifunctional networks are primarily extended networks (green bar in Figure 1D). Analysis of the mean robustness for each network class shows that Noise-amplifying networks are the most robust, followed by static Turing networks, Multifunctional networks, and Traveling waves (Figure 1E). The atlas shows a similar proportion of Type I and Type III networks, which can form static Turing patterns with differential diffusion (*d* > 1) or any diffusion value (*d* ≠ 0), and a minority of Type II networks that generate patterns for *d* < 1 (Figure 1F).

The analytical predictions were confirmed by numerical simulations shown in Figure 1F-I, details provided in Material and Methods and SI Appendix.

### Pattern Phase and Diffusion Constraints

Next, we derived an atlas with the subset of networks capable of forming static Turing patterns (yellow nodes in Figure 1B), including Mul-tifunctional networks capable of both static and oscillatory patterns (green nodes in Figure 1B), but excluding traveling waves and noise amplifiers (Figure S1).

For each node in this reduced atlas, we calculated the type of diffusion constraint and phase relationships of the network analytically. Specifically, we identified Type I networks that require different diffusion rates, Type II networks that allow equal diffusion rates, and Type III networks that allow any combination of diffusion rates. These conditions were determined based on stability conditions for homogeneous steady states and diffusion-driven instability. The phase relationships between periodic patterns formed by the network were analyzed using the relative sign of eigenvectors, reflecting four possible phases between the three reactants, as detailed in the Materials and Methods section.

The analysis revealed that in the reduced atlas, networks with similar diffusion constraints or similar phases cluster in topological space (Figure 2A), suggesting that specific regulatory feedbacks determine patterning constraints and behavior. To characterize these regulatory feedbacks, we analyzed two key transitions in the atlas, shown by boxes in Figure 2A.

**Fig. 2.**
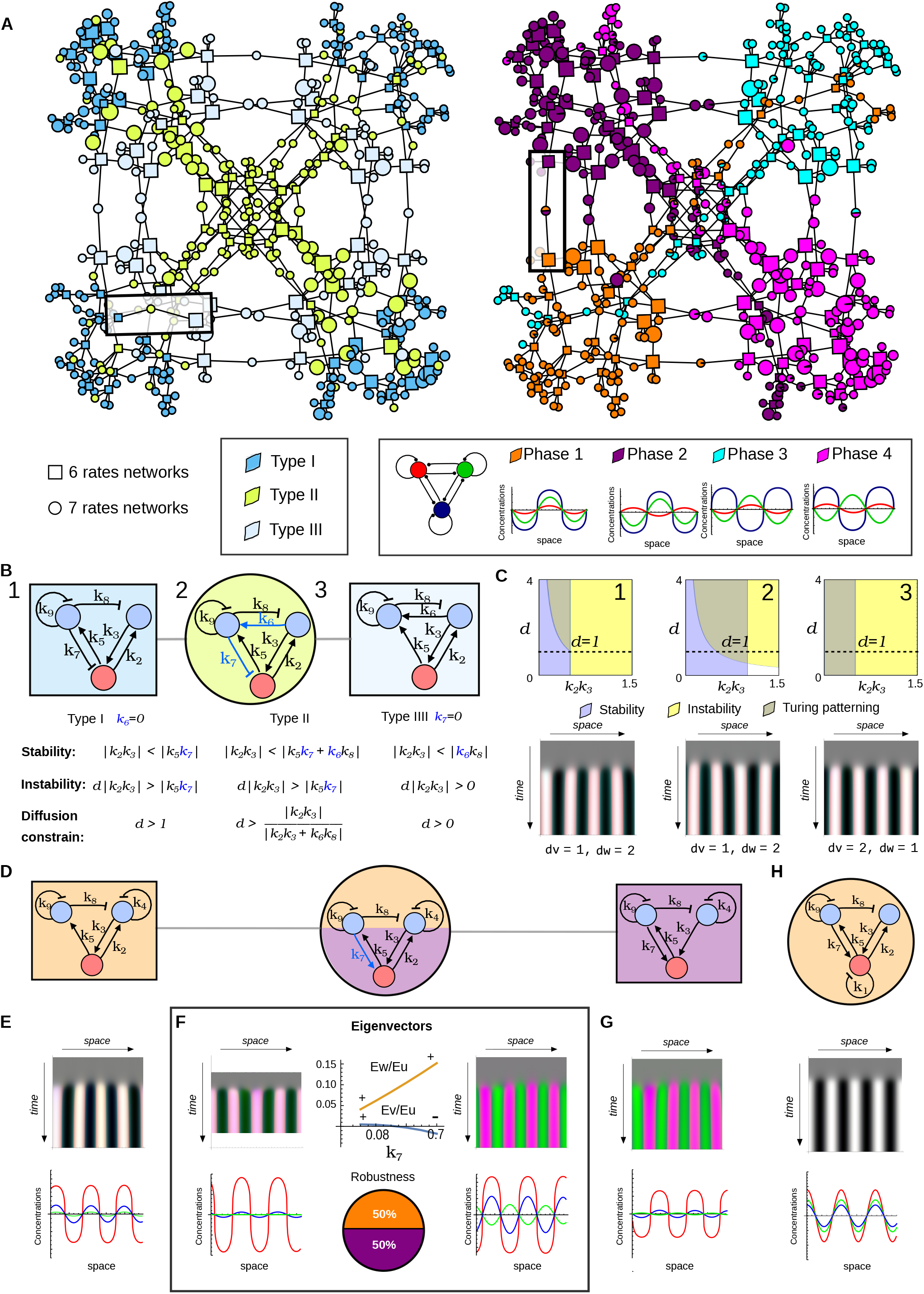
Reduced atlas of static Turing networks: topology determines diffusion constrains and pattern phase. A) A sub-graph of the atlas shown in Figure 1B obtained by considering only static Turing networks (yellow and green nodes). Left: Nodes are colored according to network type: Type I (blue) requires differential diffusivity, Type II (light green) can have equal diffusivity, Type III (light blue) allow for any diffusivity. The square highlights the transition shown in panel (B). Right: Nodes are colored according to the relative phase between the periodic patterns (legend below). The square highlights the network shown in panel D. Node size corresponds to robustness to parameter changes. B-C) Analysis of the transition between a Type I, Type II and Type III network shown in (A) on the left. B) A trade off between homogeneous steady state stability (stability) and diffusion-driven instability (instability) conditions determine the constrain on diffusion coefficient ratio d for the three networks (1,2,3). C) Top: graphs showing the parameter space that satisfy homogeneous steady state stability (blue) and diffusion driven instability (yellow), dash line shows equal diffusivity for *d* = 1. In Type II and Type III networks the the yellow and blue region intersect for *d* < 1. Bottom: Simulation of the networks (1,2,3) with *d* = 2, *d* = 1, and *d* = 0.5 respectively. D-G) Analysis of the Transition between a Phase1, Phase1/3 and Phase3 networks shown in (A) on the right, from left to right the transition is made by adding the interaction *k*_7_ and loosing interaction *k*_2_. E) Space time plot (top) and graph (below) show that the 6 edges network shown on the left in (D) forms periodic patterns of the three reactants (u,v,w) that are in phase (Phase 1). F) The 7 edges network show in the center in (D) can form periodic patterns with two different phase depending on the strength *k*_7_. Left: for low values of *k*_7_ the patterns are in phase (Phase 1), Right: for high values of *k*_7_ the patterns of u,w (red,blue) in phase but v (green) out of phase (Phase 3) as reflected by the relative sign of the Eigen vectors (Eu,Ev,Ew). Pie chart shows that Phase 3 is more robust than Phase 1 since it is formed for 64% of the parameter space. G) The 6 edges network shown on the right in (d) forms periodic patterns Phase 3. H) Example of a 7 edges network that forms only patterns in Phase 1.

The first transition considered a Type I, a Type II, and eventually a Type III network (Figure 2B). The atlas showed that transitioning between a Type I and a Type III network always requires passing through a 7-edge Type II network. In the specific transition considered, the analytical conditions for homogeneous steady-state stability and diffusion-driven instability (Figure 2B) showed that adding a new interaction *k*_6_ in the intermediate network introduces a negative feedback that enlarges the homogeneous steady-state stability parameter space, transitioning from a Type I to a Type II network allowing for *d* ≤ 1 (Figure 2C middle). Conversely, removing an interaction involved in the stability feedback (i.e., *k*_7_) enlarges the diffusion-driven instability space, making the network unstable for any *d* (Figure 2C right), transitioning from a Type II to a Type III network.

The second transition we studied was from a Phase 1 to Phase 1/3 and eventually to a Phase 3 network (Figure 2F). The atlas revealed that transitioning between Phase 1 and Phase 3, or between Phase 2 and Phase 4, always requires passing through a Multiphase network capable of both phases. In the specific transition considered, passing from a Phase 1 to a Multiphase 1/3 network involved adding the interaction *k*_7_, which introduces a new positive feedback. Altering the strength of this feedback by increasing *k*_7_ could change the relative sign of one eigenvector, altering the phase relationships of one of the reactants (Figure 2F). This demonstrated that the relative strength between the destabilizing positive feedbacks in the network determines the phase of the network. The pie chart in Figure 2F shows the parameter space percentages for each phase in the Multiphase network. We also observed that not all phase transitions are possible due to specific topological constraints, as Multiphase networks are not found between all phase pairs. Specifically, no direct transition through an intermediate Multiphase network is possible between Phase 1 and 2, and between Phase 3 and 4 (Figure 2A).

Finally, we also studied neutral transitions in the reduced atlas, where the addition of interactions did not change the relative pattern phase. Specifically, we explored transitions among networks that generate in-phase patterns to investigate the possible evolutionary trajectory of the Nodal-Lefty system (30), as shown in Figure S2.

### Compressed Topological Atlas and Regulatory Logic

Previously, we showed that any Turing network can be partitioned into network cycles, with 3-node networks having a maximum of eight cycles (*c*_1_ to *c*_8_, Figure 3A) (23, 25). We also demonstrated that Turing instability conditions can be rewritten in terms of network cycle weights and their signs (23, 25). Furthermore, networks represented by a set of cycle signs correspond to various network topologies (i.e., sets of rates *k*_1..9_), each generating distinct phases of periodic patterns but operating according to the same cycle weight sign logic (Figure 3B) (23, 25). This property allows us to compress the atlas shown in Figure 1B into a smaller atlas, where multiple network topologies map to a single network represented in terms of cycles (Figure 3B-C). In this compressed atlas, connected nodes still represent networks differing by one interaction, introducing a new cycle, mirroring the structure of the larger atlas in Figure 1B.

**Fig. 3.**
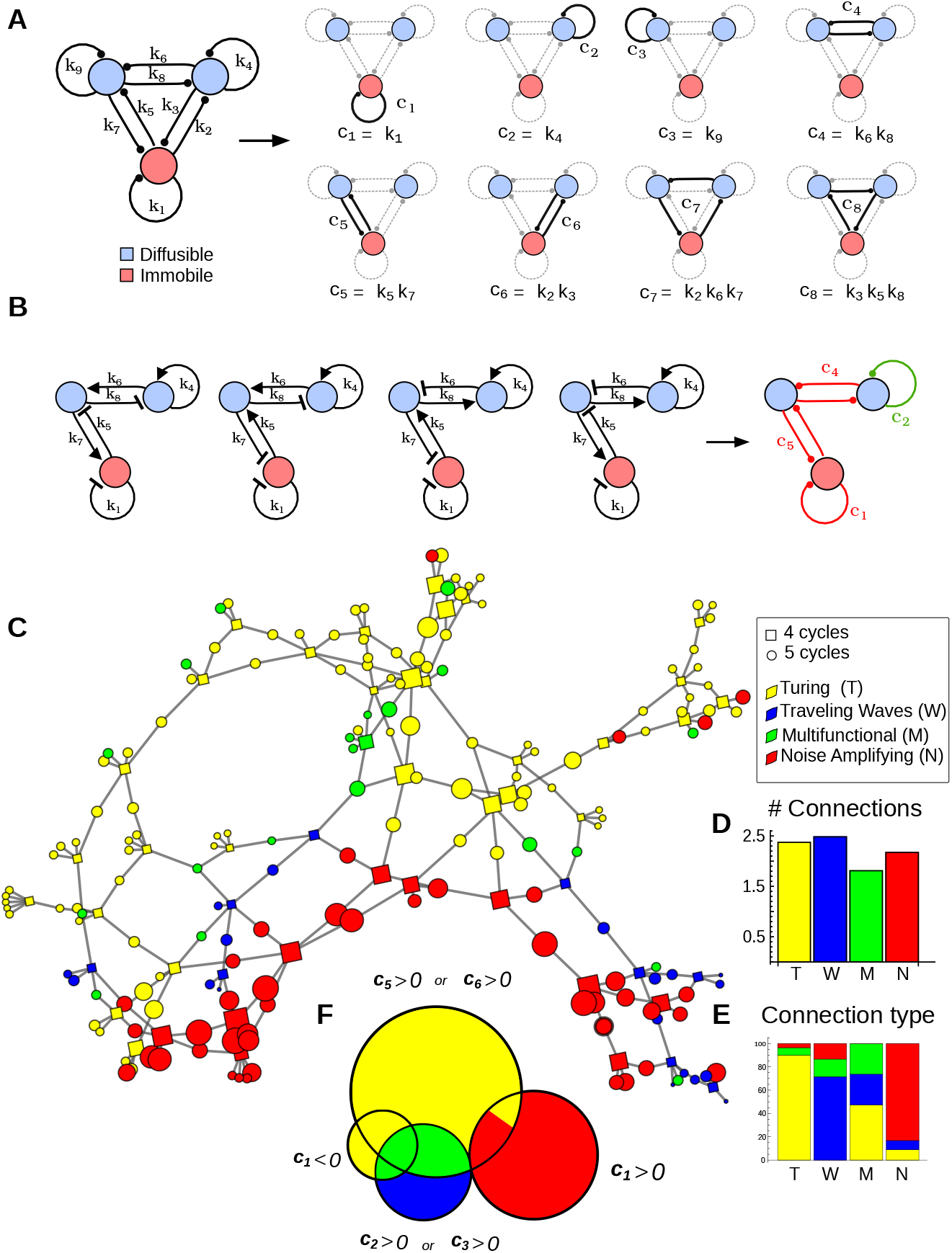
Compressed topological atlas with cycle analysis. A) The different cycles (*c*_1..8_) in a three-node gene network used to analyze Turing patterning in terms of network feedbacks. B) Cycles can be used to map multiple networks to the same underlying regulatory logic (same cycle signs). The example shows four different networks (k1..k7) mapped to a single network with negative cycles *c*_1_, *c*_4_, and *c*_5_ (red) and a positive *c*_1_ cycle (green). C) A compressed version of the atlas in Figure 1B where each node corresponds to a set of network cycles. Square nodes contain 4 cycles (6 interactions) and are connected to circular nodes with 5 cycles (7 interactions). Network colors correspond to Turing patterning behaviors: Turing (yellow, T), Traveling Waves (blue, W), Multifunctional (green, M), and Noise-Amplification (red, N). D) Average number of direct neighboring nodes per network type: Turing, Traveling Waves, and Noise-Amplifying networks have approximately 2.5 neighbors, while Multifunctional networks have approximately 1.7 neighbors. E) Proportion of neighbor types per network type: Turing nodes connect to other Turing nodes (yellow) and some Multifunctional (green) and Noise-Amplifying (red) nodes, but not to Traveling Waves (blue); Traveling Wave nodes connect primarily to other Traveling Waves (blue), Multifunctional (green), and Noise-Amplifying (red) nodes, but not to Turing networks (yellow). Multifunctional networks always mediate transitions between Turing and Traveling Wave nodes (see also panel C). F) Each circle represents networks destabilized by a specific cycle to form diffusion-driven instability. The large yellow circle corresponds to networks destabilized by *c*_5_ > 0 or *c*_6_ > 0, promoting static Turing patterns (yellow). The blue circles correspond to networks destabilized by *c*_2_ > 0 or *c*_3_ > 0, promoting Traveling Waves (blue). The red circle corresponds to networks destabilized by *c*_1_ > 0, promoting Noise-Amplifying patterns (red). The small yellow circle corresponds to networks where *c*_1_ < 0 contributes to destabilization, promoting static Turing patterns (yellow). Intersections between circles represent networks destabilized by two cycle types: green regions are Multifunctional networks destabilized simultaneously by (*c*_5_ > 0 or *c*_6_ > 0) and (*c*_2_ > 0 or *c*_3_ > 0). The yellow region between the yellow and red circles represents networks destabilized simultaneously by (*c*_5_ > 0 or *c*_6_ > 0) and (*c*_1_ > 0), promoting static Turing patterns.

As in the larger atlas, each node in the compressed atlas can still be classified into one of the behavioral classes: Turing, Traveling Waves, Multifunctional (Turing and Traveling Waves), and Noise Amplification (Figure 3C). This classification is done by studying the change of signs of the characteristic polynomial’s coefficients *a*_1_, *a*_2_, *a*_3_ promoted by diffusion-driven instability but written in terms of cycles *c*_1_ to *c*_8_ as presented in (25). This confirms the results in (25), showing that network cycles and their signs are the main determinants of the patterning capabilities of Turing networks and diffusion constrains, see also Figure S4 and SI Appendix. It also challenges previous accounts suggesting differential robustness between AIJT and CAIJT topologies (26), as we show that in linear models, these topologies have equivalent parameter space associated with the same underlying regulatory logic.

In agreement with the findings of the larger atlas in Figure 1B, we find that networks of the same type cluster together. Additionally, the compressed atlas reveals that Turing, Traveling Wave, and Noise-Amplifying networks have a similar average number of neighboring nodes (around 2.25), while Multifunctional networks have fewer connections (around 1.75), suggesting these networks may mediate connections between clusters (Figure 3D). Analysis of the neighboring node types shows that Turing networks primarily connect to other Turing networks and some Multifunctional or Noise-Amplifying networks but not to Traveling Waves. Conversely, Traveling Wave networks connect primarily to other Traveling Wave and Multifunctional or Noise-Amplifying networks but not to Turing networks. This confirms that Multifunctional networks are primarily implemented by extended networks that serve as intermediaries, connecting static Turing and Traveling Wave networks (Figure 3C). An exception is the only minimal Mutlifunctional network found in the compressed atlas, which is positioned at the center of a small cluster of Multifunctional networks.

By analyzing the condition for diffusion-driven instability derived in (25), we identified which cycles promote Turing instability for each network (Figure 3F-G, see also SI Appendix). Our analysis shows that in static Turing networks, it is primarely a positive cycle of length two between a diffusible and a non-diffusible node (*c*_5_ or *c*_6_). For traveling wave formation, it is a positive cycle of length one on a diffusible node (*c*_3_ or *c*_2_). This differs from (27), which suggested that self-regulatory positive feedback on diffusing nodes are prevalent in static Turing networks. We find that Multifunctional networks, capable of forming both Turing and traveling waves, contain both types of positive cycles. Finally, noise-amplifying networks have a positive autoregulatory feedback on the immobile node (*c*_1_), confirming that destabilizing feedback encompassing only immobile nodes is required for an asymptotic dispersion relation (23, 25).

In the next section, we show that the compressed atlas provides a comprehensive framework for understanding how modulating regulatory feedbacks in self-organizing Turing networks can drive transitions between different patterning behaviors.

### Transitions between Static Turing Patterns and Traveling Waves

To better understand how different network feedbacks control Turing patterning behaviors, we explored three trajectories in the compressed atlas.

Since our analysis of the compressed atlas (Figure 3F) showed that static Turing patterns are promoted by positive cycles *c*_5_ or *c*_6_, which can make *a*_3_ negative, while Traveling Waves networks are characterized by positive cycles *c*_2_ or *c*_3_, which can make *a*_1_ or *a*_2_ negative, we first explored a trajectory involving the gain or loss of cycles *c*_2_ and *c*_6_.

We chose the trajectory involving three nodes shown in Figure 4A-B. For each node, we performed symbolic linear stability analysis to study how changes in network cycles promoted transitions from static Turing patterns to Traveling Waves. Our analysis revealed that adding a positive cycle *c*_2_ to a network that contains a positive *c*_6_ transitions it to a Multifunctional network allowing both positive real and complex roots, shown in the bifurcation diagram in Figure 4C. Removing cycle *c*_6_ while retaining *c*_2_ further transitions the network to a minimal network capable of generating only Traveling Waves (Figure 4B).

**Fig. 4.**
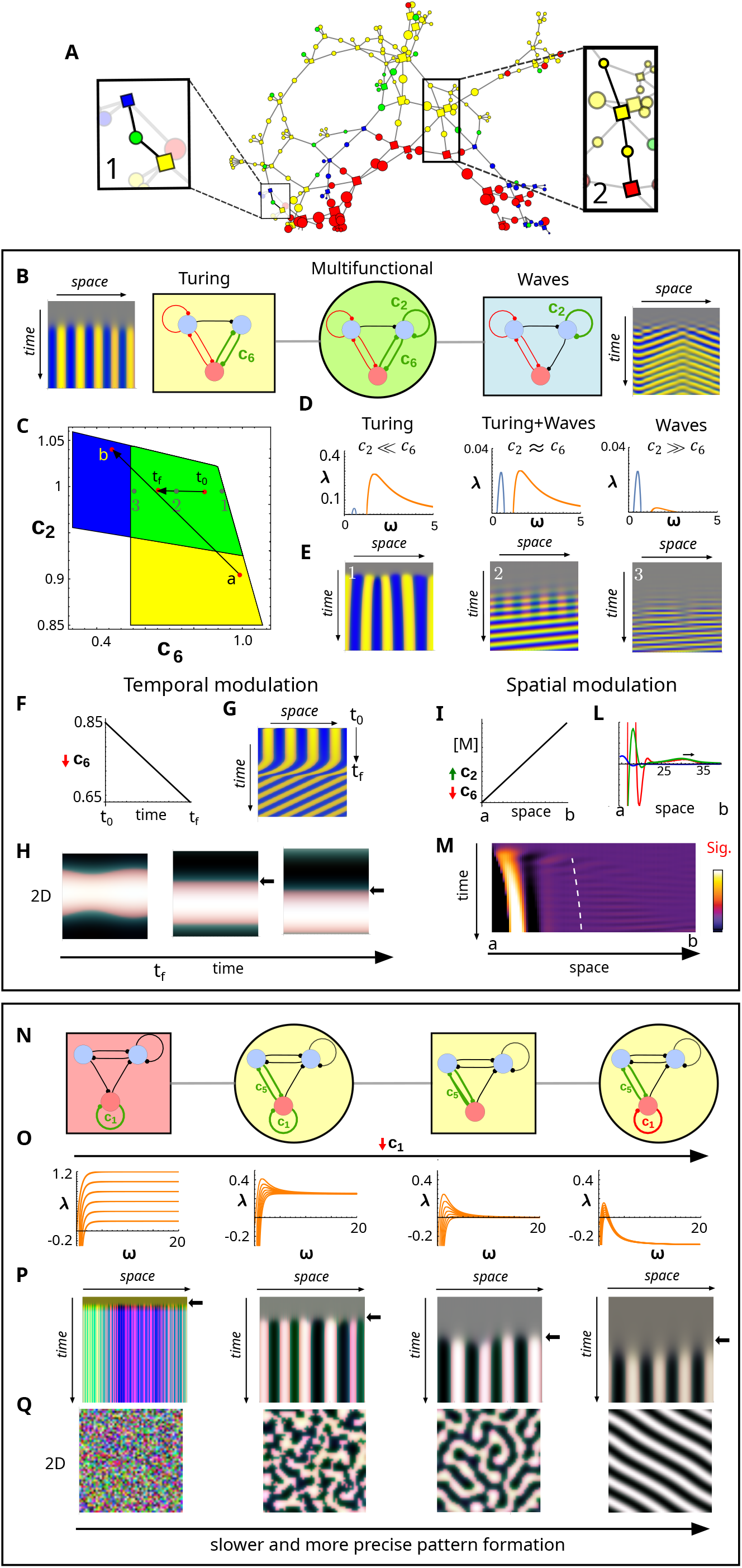
Transition between Noise amplifying, Turing and Traveling Wave networks. A) The boxes highlight two transitions analyzed in detail: 1) Transition between a Turing network (yellow square), a multifunctional network (green circle), and a traveling waves network (blue square), studied in panels B-M. 2) Transition between a noise amplifying network (red square) and static Turing networks (yellow square and circles), studied in panels N-Q. B) Details of transition 1. Left: 1D Turing network (yellow square) with a destabilizing positive cycle *c*_6_ generates a periodic static pattern (straight lines in spacetime plot). Middle: addition of new cycle *c*_2_ promotes transition from a static Turing network to a multifunctional network (green circle). Right: losing cycle *c*_6_ promotes a transition to traveling waves driven by *c*_2_ (diagonal lines in space-time plot). C) Parameter space associated with diffusion-driven instability for different strengths of *c*_2_ and *c*_6_ in the multifunctional network (green circle, panel B). The yellow region has only a positive real eigenvalue (Turing). The blue region has only a positive complex eigenvalue (Traveling Waves). The green region has both a complex and real positive eigenvalue (Turing and Waves) with points (1..3) showing parameters used in D-E, and horizontal arrow between 3 and 1 showing temporal modulation of *c*_6_ (t0 to tf) simulated in panels G-H. The arrow from point a to b shows spatial modulation used for simulations in panels L-M. D) Positive real (orange) and complex (blue) eigenvalues for parameter sets (1..3). For *c*_2_ ≪ *c*_6_, the positive real eigenvalue associated with Turing patterns is larger; for *c*_2_ ≈ *c*_6_, both eigenvalues have similar maximums; for *c*_2_ ≫ *c*_6_, the complex positive eigenvalue associated with traveling waves is larger. E) Space-time plot of simulations for parameter sets (1..3) shows the relative magnitude of the eigenvalues correctly predicts the patterning outcome. When they have equal magnitude (*c*_2_ ≈ *c*_6_), both static and traveling waves coexist. F-G) A continuous linear decrease of *c*_6_ over time (from *t*_0_ to *t*_*f*_) promotes the formation of a static Turing pattern that transforms into traveling waves (straight lines transform into diagonals in space-time plot). H) In a 2D simulation, temporal *c*_6_ modulation promotes the formation of a stripe that begins to propagate. I-L) A linear change of *c*_6_ and *c*_2_ in space (from point a to b) promoted by a morphogen M drives a transition in space from static Turing pattern to traveling waves. M) This type of transition can help interpret the self-organizing patterning dynamics observed in Gastruloids (31). N) Details of transition 2: From left to right, a decreasing *c*_1_ promotes transitions from a noise amplifying network (*c*_1_ ≫ 0, red square) to three Turing networks (*c*_1_ > 0, yellow circle; *c*_1_ = 0, yellow square; and *c*_1_ < 0, yellow circle). O) Dispersion relations for networks in panel N with decreasing values of *c*_1_ and varying *c*_5_. Ordered from left to right: noise amplifying networks with 0 < *c*_1_ < 1.2; Turing network with *c*_1_ = 0.25 and 0 < *c*_5_ < 1.19; Turing network with *c*_1_ = 0 and 0 < *c*_5_ < 1.5; and Turing network with *c*_1_ = −0.3 and 1.5 < *c*_5_ < 1.875. In the noise amplifying case, the value of *c*_1_ corresponds to the asymptote of the dispersion relation. In the other cases, as *c*_1_ decreases, the dispersion relations shift towards negative values. P) 1D simulations show that as *c*_1_ decreases, lower eigenvalues promote slower pattern formation (black cross marks later pattern appearance in space-time plots). Q) 2D simulations show that as *c*_1_ decreases, the smaller range of modes that become unstable in practice due to the eigenvalue shifting towards negative values promotes more precise two-dimensional patterns with a characteristic wavelength.

In the Multifunctional network containing both *c*_6_ and *c*_2_, the relative strength of these cycles controls the maximum values of the positive real and complex eigenvalues, determining the dominant behavior (Figure 4D). When *c*_2_ ≪ c_6_, static Turing patterns dominate; conversely, when *c*_2_ ≫ c_6_, traveling waves dominate. Numerical simulations with three parameter sets where both complex and real positive eigenvalues exist (points 1, 2, 3 in Figure 4C) confirmed that the dominant patterning behavior could be predicted by the relative maximum magnitude of the real and complex eigenvalues (Figure 4E).

To further test the predictive power of the bifurcation diagram shown in Figure 4C, we defined a trajectory within the multifunctional parameter space (green region in Figure 4C) to modulate network behavior over time (from *t*_0_ to *t*_*f*_), investigating the potential for transitioning between patterning behaviors during embryonic development. By promoting a linear reduction in *c*_6_ strength over time (Figure 4F), the Multifunctional network first formed a static Turing pattern that then transformed into Traveling waves, highlighted by a space-time plot in Figure 4G. In a 2D domain slightly smaller than the wavelength, this modulation generated a straight stripe that propagated along its axis (Figure 4H).

Finally, to further explore the ability of the Multifunctional network, we defined a modulation over a wider range of the parameter space by decreasing *c*_6_ and increasing *c*_2_ simultaneously, see the arrow from point a to b in Figure 4C. This modulation transitioned from a pure Turing region (a), having only a positive real eigenvalue, to a pure traveling wave region (b), having only a positive complex eigenvalue. Our aim was to test whether this wider modulation could be promoted by a linear morphogen gradient M, driving a transition between static Turing patterns to traveling waves over space rather than time (Figure 4I). In agreement with our predictions, one-dimensional simulations generated one peak of a static Turing pattern on one side of the spatial domain connected to a region of traveling waves on the opposite side (Figure 4L-M). This type of modulation could be relevant for several self-organizing systems such as Gastruloids (31), where the modulation of different signaling pathway feedback could be linked to the formation of an axis at one extreme of the aggregate, transitioning into traveling waves that resemble somitogenesis at the other aggregate extreme, see Figure S3.

We also analyzed a different type of transition between a Traveling Wave and a Multifunctional network, highlighted in Figure S5. As mentioned above, Static Turing Wave networks are characterized by a positive destabilizing cycle *c*_5_ or *c*_6_ that makes *a*_3_ negative. Our analysis showed that an alternative transition to Multifunctional networks can be achieved by introducing a negative cycle *c*_1_ rather than *c*_6_, which changes the contribution of *c*_2_ in the characteristic polynomial coefficient *a*_3_ making it negative (Figure 5C). This demonstrates an alternative path to Multifunctional behavior, highlighting the complex interaction between feedbacks in Turing networks to transition between patterning behaviors.

### Transition Between Turing and Noise Amplifying Networks

In the atlas, both static Turing networks and Noise-amplifying networks are predominant (Figure 1B), and direct transitions are possible between these two network types (Figure 3C). As shown in Figure 3F, most static Turing patterns are driven by cycles *c*_5_ > 0 or *c*_6_ > 0, while all Noise Amplifying networks have *c*_1_ > 0. At the intersection of these clusters, networks sometimes display normal Turing patterning behavior and sometimes noise amplification. To better understand these intermediate cases, we analyzed a transition in the compressed atlas from a noise-amplifying network to three subsequent Turing nodes, marked by changes in *c*_1_ and gains in *c*_6_, highlighted by the box on the right in Figure 4A and in Figure 4O.

In this transition, we observed that the sign and strength of *c*_1_ significantly affect the shape of the dispersion relation and pattern formation capabilities (Figure 4N-O). A positive *c*_1_ results in a higher asymptote for large wavenumbers, leading to greater unspecific amplification of modes. Lowering *c*_1_ reduces this asymptote, decreasing unspecific amplification and resulting in more precise patterns.

In agreement with this prediction, 1D and 2D simulations show that a positive *c*_1_ promotes fast patterning with random noise amplification (Figure 4P-Q). When *c*_1_ strength is smaller and coexists with *c*_6_, it leads to static Turing patterns that are noisy and have irregular wavelengths in 2D. As *c*_1_ decreases to zero, patterning slows and becomes more regular. Further negative values of *c*_1_ continue to slow down patterning and improve its regularity.

The intuitive interpretation of these results is that the self-regulatory loop *c*_1_ on the immobile node plays a crucial role in destabilizing the network because this node is not subjected to the equilibrating force of diffusion. A positive feedback on *c*_1_ enhances this destabilization, while negative feedbacks mitigate it. From a biological and evolutionary standpoint, these findings suggest that cell autonomous feedback mechanisms involving immobile nodes are essential for modulating the trade-off between the speed and precision of pattern formation.

## Discussion

The topological atlas presented in this study offers a novel framework to understand how Turing networks transition between different self-organizing behaviours in multi-cellular systems. Employing an automated algebraic method, we identified the networks that give rise to static, oscillatory, and noise-amplifying Turing patterns. This is crucial for understanding self-organization during development, where modulations of gene regulatory networks drive changes in patterning behaviour over time or space. We systematically mapped and categorized Turing networks based on their topology and patterning capabilities, advancing previous network screenings that focused primarily on networks that generate static Turing patterns (23, 24, 26, 27).

A notable feature in the atlas is that minimal networks (6 interactions) with different behaviours are connected by extended networks (7 interactions) exhibiting hybrid multifunctional behaviour. These intermediate networks can display two types of self-organizing behaviours depending on interaction strength.

This include multiphase networks that can generate periodic Turing patterns with different phase relations between reactants depending on the parameters. These networks have not been described previously and may explain changes in the relative phase of periodic patterns observed for self-organizing patterning during embryionic development. For instance, they could account for the switch between in-phase and out-of-phase patterns of BMP signaling (pSmad) and Sox9 observed in digit patterning, as shown in the supplementary material of (6).

On the other hand, we used the atlas to explore topological changes that are neutral from a patterning phase perspective but lead to more robust Turing networks. The atlas confirmed our previous proposition (23) that changes in network topology can enhance the robustness of the Nodal-Lefty system, where Nodal and its inhibitor Lefty are co-expressed and proposed to be part of a Turing network (30). Specifically, we identified a transition from a network where Lefty inhibits only the receptor to an extended network where Lefty also directly inhibits Nodal (Figure S2). This introduces a redundant interaction and improves the network’s robustness to parameter changes. Similar neutral transitions in the atlas could be exploited to study the possible implementation of other self-organizing Turing systems during developmental.

Exploiting the representation of Turing conditions in terms of cycles, as introduced in (25), we derived a compressed atlas, where several networks with the same underlying regulatory logic are mapped into one network with a set regulatory cycle signs (see Figure 3B). This allowed us to identify which regulatory modules (cycles signs) drive different Turing behaviours (Figure 3F). This is significant because defining modules within Turing networks is particularly challenging due to the extensive feedback loops characteristic of these networks, where every gene seems to be connected with every other gene. It could also provide a way to reduce larger Turing networks into equivalent smaller ones, as done in (32). The cycle decomposition approach, however, extends beyond Turing systems and can be applied to any stability analysis of PDE systems with feedback.

The cycle-based atlas also confirmed that networks with different Turing behaviours are connected by extended Multifunctional networks. For instance, networks that form static Turing patterns and those that form travelling waves are connected by networks possessing two positive feedback cycles that can promote both real and complex positive eigenvalue, as shown in Figure 4B-C. Changing the strength of these cycles alters the relative magnitude between the two eigenvalues, leading to a smooth transition between static Turing patterns and oscillations, see Figure 4D-E.

An intriguing property of these Multifunctional networks is their ability to transition between behaviour over time (Figure 4F-G) promoting scenarios where the output of one self-organizing regime acts as initial conditions for the next, implementing developmental patterning dynamics driven by stigmergy, as proposed by Sasai Yoshiki (33). This can lead to more controlled self-organizing dynamics. An example is the 2D simulation shown in Figure 4H, where a multifunctional Turing network first forms a straight stripe from noise, in the regime of a static Turing pattern on a domain sized approximately as the wavelength, which then transforms into a series of travelling waves moving along the stripe’s direction. Modulating the cycle strength to change self-organizing behaviour can also be relevant to investigating transitions between self-organizing regimes across tissues (Figure 4I-M), as seen in the Gastruloids presented in (31) (Figure S3).

Finally, the cycle-based atlas highlights that self-regulatory feedbacks on non-diffusible nodes play a critical role in controlling the stochasticity of pattern formation. Positive feedback on these nodes accelerates pattern formation but introduces more noise, while negative feedbacks slow down pattern formation and enhance precision, as shown in Figure 4N-Q. This reveals a fundamental mechanism by which developmental systems can balance the trade-off between speed and accuracy in pattern formation, a common challenge in many complex systems. It also highlights the mechanisms by which reaction-diffusion systems generate patterns by amplifying and filtering the periodic modes present in the initial noise of the system. Faster amplification and less filtering result in noisier patterns, while slower amplification and narrower filtering (e.g., narrow dispersion relation) produce more precise patterns.

Overall, the atlas can be used to interpret how Turing networks have evolved, providing a design space of Turing networks that evolution may have explored to reach specific configurations. Trajectories along the atlas represent possible pathways of topological changes that maintain Turing behaviour, associated with single regulatory changes, and can sequentially move from one patterning behaviour to another. Our calculation of robustness to parameter changes for each network indicates the likelihood of these new networks being found at random.

Additionally, the atlas, though composed of different networks, can be seen as a general map of the behaviour of a large Turing network with many feedbacks. Many biological networks, especially gene regulatory networks controlling development, possess several feedbacks. In this context, the atlas highlights the feedbacks that are more important for specific behaviours in a fully connected 3 gene regulatory network. This suggests possible directions of change that can be promoted by feedback modulation in a network during development. Although changes are continuous in the atlas thanks to the presence of multifunctional states, not all paths are allowed.

Ultimately, our approach reframes Turing’s original idea within a network-based framework, moving beyond the traditional chemical reaction perspective (1) towards a “network basis of pattern formation”. This is a step forward to relate Turing systems with the gene networks driving self-organizing patterning during development (3). It also reveals novel network designs that can help to construct complex synthetic networks capable of transitioning between different self-organizing behaviours. Recent efforts have successfully engineered a Turing network with a 3 node regulatory logic in bacteria, achieving static patterns and demonstrating the potential for further advancements towards multifunctional capabilities (34).

In the future, we believe that our approach can be expanded to include all seven diffusion-driven instability behaviours proposed by Turing and incorporate larger networks. Given the exponential increase in analytical complexity in these systems, it would be necessary to complement our current analytical approach with numerical analysis, as done in (26, 27).

## Materials and Methods

Previous studies have already proposed mathematical theorems to derive simpler analytical conditions for diffusion-driven instability in general three-reactant reaction-diffusion systems (17, 20). To simplify the conditions even further we perform a systematic automated analysis by focusing only on minimal (6 edges) and extended (7 edges) networks of three reactants, with 3 and 2 elements set to zero in the Jacobian matrix. In addition, we consider only the case where one of the three species is immobile (one element in the diagonal diffusion matrix set to zero), simplifying the conditions and broadening the criteria for diffusion-driven instability (14, 18, 23). Finally, we derived necessary and sufficient conditions for static Turing patterns and sufficient conditions for oscillatory Turing patterns by combining the Routh-Hurwitz criterion with simpler condition for the stability of three reactant system derived in (35, 36), as introduced in the supplementary material in (25). For simulations, we construct simple PDE systems from the Jacobians to simulate spatial patterning under different parameter regimes identified by the linear stability analysis.

### Conditions for Stationary and Oscillatory Turing Patterns

We derived the necessary and sufficient conditions for the formation of Turing instability by analyzing the roots of the characteristic polynomial *P* (λ) = λ^3^ +*a*_1_(*q*)λ^2^ +*a*_2_(*q*)λ+*a*_3_(*q*) obtained by linear stability analysis for each network, where λ is the eigenvalue and *q* the wavenumber. The solution to the characteristic polynomial as a function of *q* is called dispersion relation. For a diffusion-driven instability to occur, the characteristic polynomial must have all negative roots for *q* = 0 (condition 1) and at least one root with a positive real part for *q* > 0 (condition 2 or 3). Necessary and sufficient conditions for the existence of a all negative roots can be derived with the Routh-Hurwitz criterion (37), as outlined in the supplementary material of (25).

The Routh-Hurwitz criterion is obtained by constructing the Hurwitz matrix *H*, which for a polynomial of the third degree is defined as:

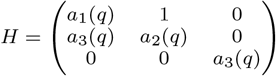

The criterion states that all roots of the polynomial have negative real parts if and only if the determinants of the leading principal minors of *H* are positive:

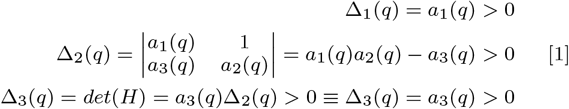

Conversely, if any of the Hurwitz terms Δ_1_, Δ_2_ or Δ_3_ becomes negative, the characteristic polynomial has at least one root with a positive real part. The number of roots with a positive real part in this case can further be estimated by the sign changes in the first column of the Routh array (37), which for a polynomial of degree three can be constructed as follows:

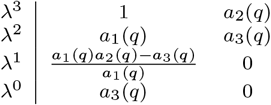

The first column of the Routh array is:

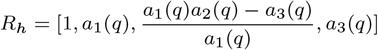

### Stationary Turing networks

Stationary Turing patterns occur when a single real eigenvalue becomes positive, specifically when the first column of the Routh array *R*_*h*_ exhibits a single sign change *R*_*h*_ = [+, +, +, −] for *q* > 0. This is a sufficient condition not only for the existence of a positive real root but also to guarantee that there are no other real positive roots. In section 2 of the supporting information, we demonstrate that for a third-degree polynomial this condition can be further simplified into:

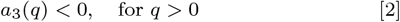

which ensures that *a*_1_(*q*) > 0 & *a*_1_(*q*)*a*_2_(*q*) − a_3_(q) > 0 guaranteeing that *R*_*h*_ = [+, +, +, −].

This represents a significant simplification since analyzing all the terms in the Routh array becomes analytically impracticable in many cases.

### Oscillatory Turing networks

Oscillatory Turing patterns occur when the characteristic polynomial has a complex positive root. This is the case when *R*_*h*_ exhibits two sign changes, *R*_*h*_ = [+, −, +, +] or *R*_*h*_ = [+, +, −, +] or *R*_*h*_ = [+, −, −, +] for *q* > 0, associated with two complex conjugate roots with a positive real part. As mentioned above, analyzing all the terms in the Routh array is often analytically impracticable. Fortunately, in section 2 of the supporting information, we demonstrate that for a third-degree polynomial this condition can be further simplified into:

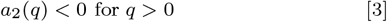

This condition simplifies the analysis considerably by guaranteeing two sign changes in the Routh array, providing necessary and sufficient conditions for the formation of oscillatory Turing patterns.

### Multifuctional Turing Networks

To identify multifunctional networks capable of both oscillatory and static patterns, we require that the simplified conditions 2 and 3 can be satisfied by the network, both independently or simultaneously. The first case identifies the parameters that give rise to either a positive real eigenvalue or a positive complex eigenvalue. The second case identifies parameters that give rise to eigenvalue that can simultaneously give rise to both situation simultenously for different wave numbers *q* (e.g., the green parameter space in Figure 4C). This allows us to pinpoint networks that can switch between oscillatory and static behaviours depending on parameter variations, providing a comprehensive understanding of the network’s multifunctional capabilities.

### Noise-amplifying Turing Networks

If a Turing network has an eigenvalue with a positive asymptote for *q* → ∞, a condition previously identified as necessary for the amplification of noise (23, 25), we verify whether the dispersion has a maximum above this asymptote to classify the network as noise amplifying.

This verification involves obtaining parameters that satisfy diffusion-driven instability with the *FindInstance* command in Wolfram Mathematica. In the case of multifunctional network we derive parameters for both static Turing patterning and traveling wave behaviour. Starting from these parameters we derive several parameter sets by allowing one parameter to vary from its minimum to its maximum allowed values as determined by the linear stability analysis conditions. For each parameter set, we calculate the asymptote of the dispersion relation using the function *Limit* for *q* → ∞. λ(*q*). Secondly we find the maximum eigenvlue numerically using the function *FindMaximum*.

If the maximum eigenvalue is lower than the limit for *q* → ∞ for all parameter sets of the network, we consider the network as a noise amplifying. Overall, we observe that if the limit is larger than the maximum, noise-amplifying networks consistently produce static patternss, regardless of the presence of a positive real root with a complex part. This supports our earlier finding that oscillatory noise-amplifying networks are only feasible in systems with four nodes (25).

### Numerical Simulation

To simulate a network, we obtained representative Jacobian values 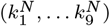 and diffusion coefficients 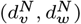 that respect patterning conditions of the network *N* with the *FindInstance* command of Wolfram Mathematica, and construct a system of PDEs for the concentration vector **c** = (*u, v, w*)^*T*^ is given by:

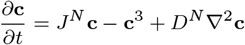

where *J*^*N*^ is the Jacobian matrix of the network obtain by substituting the parameters 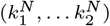, *D*^*N*^ is the diagonal diffusion matrix obtained by substituting 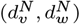, and **c**^3^ represents cubic non-linear terms that provide saturation:

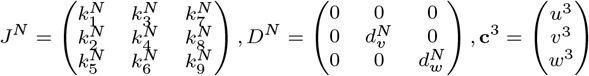

Note that *J*^*N*^ has 3 or 2 elements set to 0 in minimal and extended networks respectively, and *u* is the immobile specie with *d*_*u*_ = 0, and we consider periodic boundary conditions. This PDE systems have always one stable equilibrium at *c*_0_ = (*u*_0_, *v*_0_, *w*_0_) = (0, 0, 0) and generates periodic waves by diffusion-driven instability around this stable point. We begin the simulations with a random initial conditions for for all the three reactant uniformly distributed in the interval (−0.0005, 0.0005) around *c*_0_.

We perform 1D ans 2D simulations using a first order finite difference scheme for space discretization and and forward Euler method for time discretization, written in Wolfram Mathematica. The domain size *L* and total simulation *T* time are calculated according to the wavelength *ω* = 2Π*/q*_max_ and maximum eigenvalue λ_max_ obtained from the linear stability analysis, as follows:

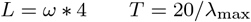

with high resolution space discretization *d*_*s*_ = *L/*300 and time discretization *d*_*t*_ = *T/*20^7^ to avoid numerical errors. All the simulations confirm as proposed by Turing (1) that the diffusion-driven behaviour can be correctly predicted by the linearized version of the system around steady state, while the non linear part *c*^3^ plays only a saturating effect for large deviation from equilibrium.

### Network Robustness Calculation

To assess the robustness of each network *N*, we quantify the volume of the parameter space that satisfies the diffusion-driven conditions derived from the linear stability analysis. This involves integrating all the diffusion-driven instability conditions *f* and it is done by fixing one negative feedback rate at −1 and one diffusion coefficient at 1, thereby calculating the relative parameter space. The relative parameter space volume is calculated over the ranges: 0.1 to 10 for reaction rates *k*_*i*_ and 0.001 to 100 for the relative diffusion coefficient ratio *d*.

The robustness *R*(*f*) is thus calculated as the integral over the defined parameter space:

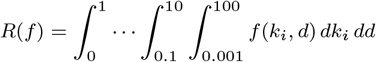

After obtaining *R*(*f*) for each network, we standardize the robustness values by dividing each *R*(*f*) by the maximum robustness value observed (i.e the most robust network), *R*(*f*_max)_. The normalized atlas robustness *r*(*f*) for each network *i* is then given by:

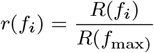

The size of a node representing a network N in the atlas is logarithmically proportional to *r*(*f*_*i*_). This approach quantifies the likelihood of a given network achieving diffusion-driven instability with randomly assigned parameters, and also provides a measure of the network’s robustness to parameter changes.

### Diffusion Constrain

As introduced in (23), Turing networks with an immobile specie exhibit distinct types of diffusion constraints for pattern-forming conditions. These constraints are categorized as follows: Type I networks require differential diffusivity, Type II networks allow equal diffusivity, and Type III networks have no specific diffusivity constraints.

The classification of network types is derived by checking weather the diffusion-driven instability conditions can be satisfied in the following cases:

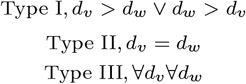

### Pattern Phase

The relative phase pattern generated by each networks, which can be categorized into four distinct configurations:

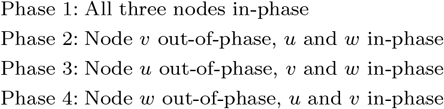

To predict the pattern phase generated by a network, we do not perform numerical simulations, instead we analyze the relative sign of the eigenvectors associated with the eigenvalue that promote diffusion-driven instability.

For a given network, we first we obtain a set of parameters (*k*_1_..*k*_9_) with the *FindInstance* command in Wolfram Mathematica that satisfy the diffusion-driven instability conditions. We let each parameter *k*_*i*_ to change within diffusion-driven instability range and calculate in each case the eigenvectors *E*(λ(*q*_max_)) = (*E*_*u*_(*q*_max_), *E*_*v*_ (*q*_max_), *E*_*w*_ (*q*_max_) associated with the positive eigenvalue values. For each case we calculate the relative signs of the eigenvectors as:

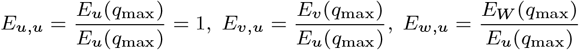

The relative sign of the phase vector *φ* = (*E*_*v,u*_, *E*_*w,u*_) determines the phase of the periodic patterns:

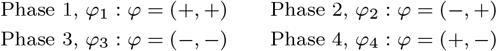

The eigenvectors can be plotted as a function of *k*_*i*_ to identify Multiphase networks, as shown in Figure 2F.

## ACKNOWLEDGMENTS

We would like to thank Xavier Diego for the useful discussions that inspired this project and Isaac Salazar-Ciudad for providing critical feedback.

