## supporting information for "The Network Basis of Pattern Formation: A Topological Atlas of Multifunctional Turing Networks"

**This PDF file includes:**

Supporting text
Figs. S1 to S5
Tables S1 to S17
SI References

### Supporting Information Text

The goal of the theoretical screening presented in this study is to analyze and relate a minimal ensemble of Turing networks that can form static and oscillatory self-organizing patterns. Turing showed that while static periodic waves can be generated by a reaction-diffusion system of 2 reactants, oscillatory behaviors such as traveling waves require at least 3 reactants (1), a finding further supported by (2). Previous studies (3, 4) have demonstrated that both static and oscillatory patterns can arise in three-reactant systems with one immobile reactant that has a zero diffusion coefficient. In our previous study (5), we also demonstrated that these are minimal Turing models exhibiting relaxation of diffusion constraints to Type II and Type III networks, allowing for equal or any combination of diffusion coefficients.

Following the principle of parsimony, also known as Ockham's Razor (6), we focus on minimal models consisting of three species with one immobile reactant, representing the smallest system capable of producing both static and oscillatory Turing patterns. This principle advocates for the simplest yet sufficient explanation (i.e., model) to understand complex phenomena. In the 14th-century words of William of Ockham:

> It is futile to do with more what can be done with fewer. Therefore, no plurality should be posited without necessity... If something can be explained by fewer causes, it is superfluous to posit more causes... By adhering to this principle, we ensure that our explanations remain both simple and effective, avoiding the introduction of unnecessary complexities.

Setting aside the contradiction that, as an English Franciscan friar, Ockham's ultimate trick was to reduce everything to a single cause of God. An idea that's undeniably parsimonious, though perhaps a bit weak on the evidence front. In biology, parsimony has often been overlooked in favor of large models with numerous variables and parameters that aim to explain the full range of biological complexity. However, we argue that minimal models are crucial for uncovering fundamental theoretical principles underlying biological systems, as demonstrated with negative feedback mechanisms to understand oscillations or feed-forward loops to interpret hierarchical signal processing. These principles have significantly advanced our understanding of how gene-regulatory networks drive different biological behaviors by focusing on simplified systems. By applying this approach to self-organization, we aim to reveal principles that can improve our understanding of diffusion-driven instabilities in gene regulatory networks.

More complex models can then be built upon these principles to relate the models with the large regulatory networks controlling embryonic development. This incremental approach ensures that models are based on well-understood principles.

### 1. Atlas construction (Figure 1)

In this section, we provide a detailed description of the construction of the topological atlas of 3-node Turing networks with one immobile node. The atlas is constructed by listing all possible Turing networks organized according to their topology. Each network is labeled (colored) according to its self-organizing behaviour.

Previously we showed that the minimal number of interactions (edges) required by 3-node network to be able to form static Turing patterns is 6 (5). Therefore we begun the construction of the atlas by screening all possible minimal Turing networks with 6 interactions.

**A. Systematic screening of minimal 3-node networks.** Given a $3 \times 3$ reaction matrix $R$ that represents all possible regulations $k_i$ between the 3 nodes in a network:

$$R = \begin{pmatrix} k_1 & k_3 & k_7 \\ k_2 & k_4 & k_8 \\ k_5 & k_6 & k_9 \end{pmatrix}$$

We define $\Omega_6$, as the set of all possible network topologies with 6 interactions obtained by setting 3 regulations $k_i$ to 0 and varying the signs of the remaining $k_i \neq 0$.

Let $C^i$ be a $3 \times 3$ matrix representing the connectivity of network $i$, containing 0s where the rates $k_i$ are set to 0 and 1s otherwise.

A reaction matrix $R^a$ of network $a$ is obtained by multiplying $R$ with $C^a$:

$$C^a = \begin{pmatrix} 1 & 1 & 1 \\ 1 & 1 & 0 \\ 1 & 0 & 0 \end{pmatrix} \quad R^a = R \circ C^a = \begin{pmatrix} k_1 & k_3 & k_7 \\ k_2 & k_4 & 0 \\ k_5 & 0 & 0 \end{pmatrix}$$

The set $\Omega_6$ of all reaction matrices $R^i$ can be calculated as the number of possible connectivity matrices $C^i$ obtained by setting 3 out of 9 elements to 0:

$$N_1 = \binom{9}{3} = \frac{9!}{3!(9-3)!} = 84$$

Next, for each reaction matrix $R^i$, we derive all possible network topologies by changing the signs of the non-zero reaction rates $k_i$. Let $T^i$ be a $3 \times 3$ matrix representing the topology of the network, where each element is 1 if the interaction is positive, -1 if it is an inhibition, and 0 otherwise.

This is achieved by multiplying $R^i$ by all possible topology matrices $T^i_j$, representing a combinations of the signs of the six non-zero rates $k_i$.

For example, a possible topology matrix for network $a$ is:

$$T^a = \begin{pmatrix} 1 & 1 & -1 \\ 1 & -1 & 0 \\ 1 & 0 & 0 \end{pmatrix}$$

The number of possible topologies $N_2$ for each network can be calculated by considering all the combinations of the signs of the six non-zero rates $k_i$ as:

$$N_2 = 2^6 = 64$$

In total, the set of possible minimal network topologies $\Omega_6$ with 6 rates is:

$$N_1 N_2 = 84 \times 64 = 5376$$

**B. Identification of Turing networks by linear stability analysis.** To identify the networks capable of diffusion-driven instability, for each network we perfom a symbolic linear stability analysis using the computer algebra capabilities of Wolfram Mathematica. We consider a system of three reactants with concentrations given by the vector $\mathbf{c} = (u, v, w)$. The dynamics of these concentrations can be described by the reaction-diffusion equations:

$$\frac{\partial \mathbf{c}}{\partial t} = \mathbf{f}(\mathbf{c}) + D\nabla^2 \mathbf{c}$$

where $\mathbf{f}(\mathbf{c})$ represents the reaction kinetics, and $D$ is the diffusion matrix. For simplicity, we assume $D$ is diagonal with the form:

$$D = \begin{pmatrix} 0 & 0 & 0 \\ 0 & d_v & 0 \\ 0 & 0 & d_w \end{pmatrix}$$

This means $u$ is immobile, while $v$ and $w$ can diffuse.

Such system can be represented by the following network diagram:

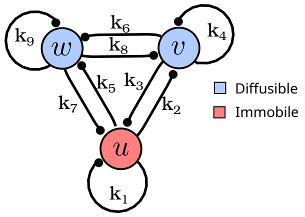

Where network interactions $k_i$, which are defined by a reaction matrix $R$, represent the elements of the Jacobian matrix $\mathbf{J}$ of $f(c)$ evaluated at the steady state $\mathbf{c}^*$:

$$R = \mathbf{J} = \left. \frac{\partial \mathbf{f}}{\partial \mathbf{c}} \right|_{\mathbf{c}=\mathbf{c}^*}$$

To analyze the stability, we introduce small perturbations around the steady state $\mathbf{c}^*$:

$$\mathbf{c}(\mathbf{x}, t) = \mathbf{c}^* + \epsilon(\mathbf{x}, t)$$

where $\epsilon(\mathbf{x}, t)$ represents small deviations from the steady state.

The perturbations $\epsilon$ can be expressed as plane waves:

$$\epsilon(\mathbf{x}, t) = \epsilon_0 e^{\lambda t + i\mathbf{q}\cdot\mathbf{x}}$$

where:
- $\epsilon_0$ is the amplitude of the perturbation.
- $\lambda$ is the growth rate (eigenvalue).
- $i$ is the imaginary unit, indicating the spatial oscillation.
- $\mathbf{q}$ is the wavevector, indicating the spatial frequency of the oscillation.
- $\mathbf{x}$ is the spatial coordinate.

This form allows us to study how perturbations evolve over time and space.

Substitute $\mathbf{c} = \mathbf{c}^* + \epsilon$ into the reaction-diffusion equations and linearize by neglecting higher-order terms:

$$\frac{\partial(\mathbf{c}^* + \epsilon)}{\partial t} = \mathbf{J}\epsilon + D\nabla^2\epsilon$$

Since $\mathbf{c}^*$ is a steady state, $\mathbf{f}(\mathbf{c}^*) = 0$, and we get:

$$\frac{\partial\epsilon}{\partial t} = \mathbf{J}\epsilon + D\nabla^2\epsilon$$

Substituting the plane wave form of $\epsilon$:

$$\frac{\partial\epsilon_0 e^{\lambda t + i\mathbf{q}\cdot\mathbf{x}}}{\partial t} = \mathbf{J}(\epsilon_0 e^{\lambda t + i\mathbf{q}\cdot\mathbf{x}}) + D\nabla^2(\epsilon_0 e^{\lambda t + i\mathbf{q}\cdot\mathbf{x}})$$

The time derivative yields:

$$\lambda\epsilon_0 e^{\lambda t + i\mathbf{q}\cdot\mathbf{x}}$$

The Laplacian term becomes:

$$\nabla^2(\epsilon_0 e^{\lambda t + i\mathbf{q}\cdot\mathbf{x}}) = -\mathbf{q}^2\epsilon_0 e^{\lambda t + i\mathbf{q}\cdot\mathbf{x}}$$

So the equation simplifies to:

$$\lambda\epsilon_0 e^{\lambda t + i\mathbf{q}\cdot\mathbf{x}} = \mathbf{J}\epsilon_0 e^{\lambda t + i\mathbf{q}\cdot\mathbf{x}} - D\mathbf{q}^2\epsilon_0 e^{\lambda t + i\mathbf{q}\cdot\mathbf{x}}$$

Canceling the exponential terms, we get:

$$\lambda\epsilon_0 = (\mathbf{J} - D\mathbf{q}^2)\epsilon_0$$

This is an eigenvalue problem. To find the eigenvalues $\lambda$, we solve the characteristic polynomial:

$$\det(\lambda\mathbf{I} - (\mathbf{J} - D\mathbf{q}^2)) = 0$$

where $\mathbf{I}$ is the identity matrix. Expanding this determinant, we obtain the characteristic polynomial in $\lambda$:

$$P(\lambda) = \det(\mathbf{J} - D\mathbf{q}^2 - \lambda\mathbf{I}) = \lambda^3 + a_1(q)\lambda^2 + a_2(q)\lambda + a_3(q) = 0$$

where the coefficients $a_1(q), a_2(q)$, and $a_3(q)$ are functions of the wavenumber $q$ and the elements of the Jacobian matrix $\mathbf{J} = \mathbf{R}$ and the diffusion matrix $\mathbf{D}$.

The solutions of characteristic polynomial $\lambda_1, \lambda_2, \lambda_3$ correspond to the eigenvalues and their signs determine the stability and pattern-forming capabilities of the system. If the real part of all eigenvalues is negative the system is stable. Conversely if at least one eiganvalue has a positive real part the system is unstable.

A standard cubic Turing system must respect two conditions:

1) Stability of the homogeneous steady state:
$\Re(\lambda_i) < 0, \quad \forall i \in [1, 2, 3]$, for $q = 0$,

2) Diffusion-driven instability:
$\exists i \in [1, 2, 3], \quad \Re(\lambda_i) > 0$, for $q > 0$

### 1. Homogeneous steady state stability condition

Homogeneous steady state stability requires the all eigenvalues $\lambda_i$ must have a negative real part for $q = 0$ to be stable. Since finding the actual analytical solution $\lambda_i$ algebraically is mathematically challenging even for small three reactant systems, an alternative way to ensure the stability of the system is to derive necessary and sufficient conditions for the negativity of all roots of the characteristic polynomial, with the Routh-Hurwitz criterion (7). This consists in checking the positivity of specific terms $\Delta_i(q)$ obtained by linear combination of the coefficients of the characteristic polynomial $a_i(q)$. For a cubic system the simple conditions for stability is defined by the Liénard-Chipart form of Routh-Hurwitz criterion:

$$\Delta_1(q) = a_1(q) > 0$$

$$\Delta_2(q) = a_1(q)a_2(q) - a_3(q) > 0$$

$$\Delta_3(q) = a_3(q) > 0$$

So to ensure stability in the absence of diffusion we check that the networks respect:

$$\triangle_1(0) > 0, \ \triangle_2(0) > 0 \text{ and } \triangle_3(0) > 0$$

### 2. Diffusion-driven instability conditions

Diffusion-driven instability requires that the real part of at least one eigenvalue $\lambda_i$ has to becomes positive for some value of $q > 0$ to make the system unstable.

To construct a atlas of static and oscillatory Turing networks, we want to further check if the diffusion-driven instability is caused by a positive root with a negative complex part, which promotes the formation of static periodic pattern, or by root with a positive complex part, which is associated with oscillatory patterns, i.e. traveling waves.

As already explained by Turing (1), the latter case is possible only with at least three reactants including the case of three reactant networks with an immobile node considered in this study. Typically, this type of oscillatory behaviour happens when two solutions have positive real parts, one with a negative complex part and the other with a positive complex part, which is known as a pair of complex conjugates. In summary, periodic static patterns require one eigenvalues to turn positive, while oscillatory patterns typically require at least two eigenvalues to turn postive.

The existence of at least a $\lambda_i$ with a positive real part is guaranteed by a violation of the Liénard-Chipart criteria:

$$\text{For } q > 0, \ \triangle_1(q) > 0 \text{ or } \triangle_2(q) < 0 \text{ or } \triangle_3(0) > 0$$

This, however, does not allow us to determine how many roots become positive. Fortunately, the number of roots with positive real parts can be estimated using a corollary of the Routh-Hurwitz criterion, which states that the maximum number of roots with a positive real part equals the number of sign changes in the first column of the Routh array (7). A similar analysis for a three-reactant Turing system with an immobile node has been presented in (4). Here, we adopt an analogous approach but further simplify the conditions to derive a set of sufficient diffusion-driven instability conditions in terms of the individual coefficients $a_i(q)$ of the characteristic polynomial. This provides a new set of minimal conditions for studying the number of roots with a positive real part in a cubic polynomial.

We begin by considering the Routh array associated with the cubic polynomial $P(\lambda)$ as follows:

$$\begin{array}{c|cc} \lambda^3 & 1 & a_2(q) \\ \lambda^2 & a_1(q) & a_3(q) \\ \lambda^1 & \frac{a_1(q)a_2(q)-a_3(q)}{a_1(q)} & 0 \\ \lambda^0 & a_3(q) & 0 \end{array}$$

We define $R_h$ as the first column of the array:

$$R_h = [1, a_1(q), \frac{a_1(q)a_2(q) - a_3(q)}{a_1(q)}, a_3(q)]$$

For sign changes, we consider a switch from + to - (or vice-versa) as we move along $R_h$. For example, no sign changes with $R_h = [+, +, +, +]$ indicate a stable system with all negative roots. One sign change, e.g., $R_h = [+, +, +, -]$, corresponds to one positive real root; and two sign changes, e.g., $R_h = [+, +, -, +]$ or $R_h = [+, -, -, +]$, correspond to two positive real roots. To derive simpler conditions, we study how the signs of $R_h$ change as we systematically vary the signs of $a_1$, $a_2$, and $a_3$, where $a_i$ is a shorthand for $a_i(q)$. In each case, we also analyze the sign of $a_1a_2 - a_3$, if needed, to determine the sign of the third element of $R_h$. The possible cases are summarized in the following table:

| Case | $a_1$ | $a_2$ | $a_3$ | $R_h$ | Discriminant | Positive Roots |
|---|---|---|---|---|---|---|
| 1. | $+$ | $+$ | $+$ | $[+,+,+,+]$ | $a_3 < a_1a_2$ | 0 |
| 2. | $+$ | $+$ | $+$ | $[+,+,-,+]$ | $a_3 > a_1a_2$ | 2 |
| 3. | $+$ | $+$ | $-$ | $[+,+,+,-]$ | $a_3 < a_1a_2$=**True** | 1 |
| 4. | $+$ | $+$ | $-$ | $[+,+,-,-]$ | $a_3 > a_1a_2$=**False** | 1 |
| 5. | $+$ | $-$ | $+$ | $[+,+,+,+]$ | $a_3 < a_1a_2$=**False** | 0 |
| 6. | $+$ | $-$ | $+$ | $[+,+,-,+]$ | $a_3 > a_1a_2$=**True** | 2 |
| 7. | $+$ | $-$ | $-$ | $[+,+,+,-]$ | $a_3 < a_1a_2$ | 1 |
| 8. | $+$ | $-$ | $-$ | $[+,+,-,-]$ | $a_3 > a_1a_2$ | 1 |
| 9. | $-$ | $+$ | $+$ | $[+,-,+,+]$ | $a_3 < a_1a_2$=**True** | 2 |
| 10. | $-$ | $+$ | $+$ | $[+,-,-,+]$ | $a_3 > a_1a_2$=**False** | 2 |
| 11. | $-$ | $+$ | $-$ | $[+,-,+,-]$ | $a_1a_2 < a_3$ | 3 |
| 12. | $-$ | $+$ | $-$ | $[+,-,-,-]$ | $a_1a_2 > a_3$ | 1 |
| 13. | $-$ | $-$ | $+$ | $[+,-,+,+]$ | $a_3 < a_1a_2$ | 2 |
| 14. | $-$ | $-$ | $+$ | $[+,-,-,+]$ | $a_3 > a_1a_2$ | 2 |
| 15. | $-$ | $-$ | $-$ | $[+,-,+,-]$ | $a_3 < a_1a_2$ = **False** | 3 |
| 16. | $-$ | $-$ | $-$ | $[+,-,-,-]$ | $a_3 > a_1a_2$=**True** | 1 |

**Table S1**

Here we can observe that four cases (4, 5, 10, and 15) in table S1 can never be satisfied because the sign of the third element of $R_h$ cannot match the specified set of signs for the coefficients $a_i$. Additionally, in some cases, the number of positive roots remains the same, regardless of the sign of the third element of $R_h$. For the purpose of deriving conditions that promote static or oscillatory diffusion-driven instability, such cases are equivalent, allowing us to merge cases (7, 8) and cases (13, 14).

Grouping equivalent cases and removing the False cases leaves us with the following reduced table where one root with a positive real part correspond to a static diffusion-driven instability, because in an oscillatory system the existence a root with a complex positive part is associated with a complex conjugate pair where two roots have a positive real part. Conversely, if at least two roots have a positive real part it is likely that one of the root has a positive complex part:

| Case | $a_1$ | $a_2$ | $a_3$ | Discriminant | Positive Roots | Probable instability |
|---|---|---|---|---|---|---|
| 1. | $+$ | $+$ | $+$ | $a_3 > a_1a_2$ | 2 | unstable oscillatory |
| 2. | $+$ | $+$ | $-$ | True | 1 | unstable static |
| 3. | $+$ | $-$ | $+$ | True | 2 | unstable oscillatory |
| 4. | $+$ | $-$ | $-$ | True | 1 | unstable static |
| 5. | $-$ | $+$ | $+$ | True | 2 | unstable oscillatory |
| 6. | $-$ | $+$ | $-$ | $a_1a_2 < a_3$ | 3 | unstable oscillatory |
| 7. | $-$ | $+$ | $-$ | $a_1a_2 > a_3$ | 1 | unstable static |
| 8. | $-$ | $-$ | $+$ | True | 2 | unstable oscillatory |
| 9. | $-$ | $-$ | $-$ | True | 1 | unstable static |

**Table S2**

To further simplify the diffusion-driven instability condition table S2, we consider the characteristic polynomial $P(\lambda)$ in its factored form, in terms of the roots $\lambda_i$:

$$P(\lambda) = (\lambda - \lambda_1)(\lambda - \lambda_2)(\lambda - \lambda_3)$$

Expanding this factored form, we obtain the polynomial:

$$P(\lambda) = \lambda^3 - (\lambda_1 + \lambda_2 + \lambda_3)\lambda^2 + (\lambda_1\lambda_2 + \lambda_2\lambda_3 + \lambda_3\lambda_1)\lambda - \lambda_1\lambda_2\lambda_3$$

Comparing this with the standard form of the characteristic polynomial, $\lambda^3 + a_1(q)\lambda^2 + a_2(q)\lambda + a_3(q) = 0$, we identify the coefficients $a_1(q)$, $a_2(q)$, and $a_3(q)$ as follows:

$$a_1(q) = -(\lambda_1 + \lambda_2 + \lambda_3)$$

represents the sum of the eigenvalues,

$$a_2(q) = \lambda_1\lambda_2 + \lambda_2\lambda_3 + \lambda_3\lambda_1$$

represents the sum of the products of pairs of eigenvalues, and

$$a_3(q) = -\lambda_1\lambda_2\lambda_3$$

represents the product of the eigenvalues.

First, we recall that that the trace of matrix is the sum of its its eigenvalues and therefore $\mathrm{Tr}(\mathbf{J} - D\mathbf{q}^2)$=$(\lambda_1 + \lambda_2 + \lambda_3)$ and therefore $a_1(q)$ in its factored form is equivalent to:

$$a_1(q) = -\mathrm{Tr}(\mathbf{J} - D\mathbf{q}^2) = -(\mathrm{Tr}(\mathbf{J}) - \mathrm{Tr}(D\mathbf{q}^2)) = -\mathrm{Tr}(\mathbf{J}) + \mathrm{Tr}(D\mathbf{q}^2)$$

We observe that if the system is stable $a_1(0) = -\mathrm{Tr}(\mathbf{J}) > 0$ to respect the Routh Hurwitz condition of homogenous steady state stability. Thus for stable Turing systems:

$$a_1(q) > 0 \text{ because } a_1(q) = a_1(0) + \mathrm{Tr}(D\mathbf{q}^2)) > 0 \text{ for } q > 0,$$
$$\text{given that } a_1(0) > 0 \text{ and the elements of diagonal matrix } D \text{ are positive } d_v > 0, d_w > 0$$

Thus none of the cases (5 to 9) in table S2 can be satisfied by systems because they require $a_1(q) < 0$, which is incompatible with homogeneous steady state stability and can be removed.

Finally, can now check if the number of positive roots $\lambda_i$ predicted in table S2 is consistent with the signs of $a_1$, $a_2$, and $a_3$ in each case. We find that condition 1 can never be satisfied. This condition is associated with two positive roots, we find that for each possible combination $(\lambda_1, \lambda_2, \lambda_3) = ([+,+,-],[+,-,+],[-,+,+])$, the following conditions derived from the signs of $a_i$ cannot be satisfied simultaneously:

$$\lambda_1 + \lambda_2 + \lambda_3 < 0 \text{ and } \lambda_2\lambda_3 + \lambda_1(\lambda_2 + \lambda_3) > 0 = \text{False}$$

Removing the case 1 and cases 4 to 9 leaves us with the following reduced table of necessary and sufficient conditions for diffusion-driven instability:

| Case | $a_1$ | $a_2$ | $a_3$ | Stability |
|---|---|---|---|---|
| 1. | + | + | − | unstable static |
| 2. | + | − | − | unstable static |
| 3. | + | − | + | unstable oscillatory |

**Table S3**

1 and 2 are simplified necessary and sufficient conditions for having only one root with a positive real part, which corresponds to a spatially unstable system (unstable static); 3 are simplified necessary and sufficient conditions for having two roots with a positive real part, corresponding to a spatio-temporally unstable system (unstable oscillatory).
This simple conditions do not require to check the discriminant term that combines $a_1$, $a_2$, and $a_3$. This is crucial for this study because in our symbolic analysis we will retain all rates $k_i$ and diffusion coefficients $d_v$, $d_w$ as symbolic parameters and even with state-of-the-art Computer Algebra Systems such as Wolfram Mathematica, deriving full conditions becomes computationally expensive.

We can further simplify these conditions by observing that unstable static systems always require $a_3 < 0$, while unstable oscillatory patterns require $a_2 < 0$, as seen in the gray cells of Table S3. This lead to the following minimal conditions for Turing diffusion-driven instability:

**Simplified Turing Patterning Conditions**

Stability for $q = 0$:
$\triangle_1(0) > 0$, $\triangle_2(0) > 0$, and $\triangle_3(0) > 0$.

Diffusion-driven instability for $q > 0$:

1. Static Turing: $a_3(q) < 0$
2. Traveling waves: $a_2(q) < 0$
3. Multifunctional: $a_2(q) < 0$ and $a_3(q) < 0$
4. Noise-amplifying: given $\Re(\lambda_i(q)) > 0$ for $q > 0$, the maximum of $\lambda_i(q)$ is a positive asymptote for $q \to \infty$

**Table S4**

Here, Multifunctional patterning conditions are defined as satisfying both $a_3(q) < 0$ and $a_2(q) < 0$ for the same parameters. It must be noted, however, this does not necessarily correspond to the case 2 in table S3. Indeed, we are just deriving condition that guarantee that $a_3(q) < 0$ and $a_2(q) < 0$ for $q > 0$, but both conditions could be satisfied for different ranges of $q$. An example is the case presented below, where $a_3(q) < 0$ and $a_2(q) < 0$ are simultaneously satisfied by the same parameters leading to the following values of $a_2$ and $a_3$ as $q$ increases:

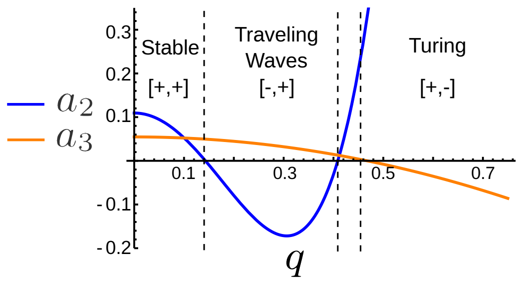

Different sign combinations of $a_2$ and $a_3$ in a Multifunctional network for parameters that respect simultaneously $a_2(q) < 0$ and $a_3(q) < 0$.

Finally, as stated in the following sections, we also consider that multifunctional networks are also those that can satisfy condition 1. and 2. in table S4 for different parameter sets.

**C. Using CAS to check Turing patterning conditions.** To derive the analytical conditions for diffusion-driven instability shown in in table S4, we use the Computer Algebra System (CAS) Wolfram Mathematica 13.

The stability conditions are obtained in terms of reaction rates $k_i$ and diffusion coefficients $d_v$ and $d_w$, by simply using the `Reduce` function of Wolfram Mathematica, which is based on Cylindrical Algebraic Decomposition.

Deriving the analytical conditions for diffusion-driven instability involves the additional complexity that the coefficients $a_i(q)$ include another positive unknown $q$. To derive conditions in terms of reaction rates ($k_i$) and diffusion coefficients $d_v$ and $d_w$, and to eliminate the unknown $q$, we follow two steps:

1. Using the `MinValue` function in Wolfram Mathematica, we derive a symbolic minimum for each coefficient with respect to $q$, denoted as $\min(a_i(q))$.

2. Using the `Reduce` function in Wolfram Mathematica, we derive negativity conditions for $\min(a_i(q))$.

The last case that we need to check is the case of noise-amplifying networks. These networks respect Turing conditions $a_2 < 0$ or $a_3 < 0$ but have a positive real eigenvalue with a positive asymptotic maximum for $q \to \infty$. The key definition here is not the asymptotic behaviour itself, but absence of a peak in the dispersion relation that would act as a filter for the growth of the possible spatial modes $q$:

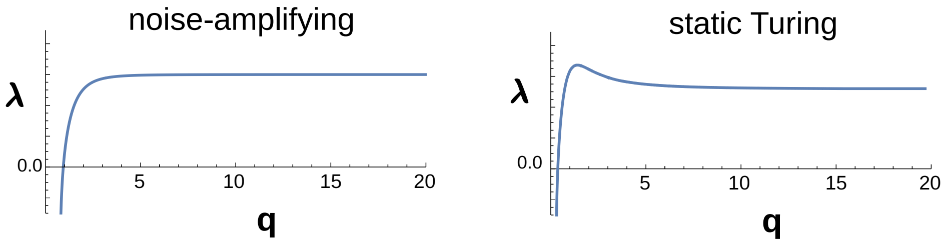

Asymptotic noise-amplifying dispersion relation (left) vs asymptotic Turing (right) with a small peak

As we previously show (5, 8) these networks, which have been recently analyzed for two reactants in (9), do not form patterns but rather amplify all modes in the initial conditions promoting noise patterns. To asses weather a network that respect the condition $a_3 < 0$ is a noise amplifying network, first we follow four steps:

1. We obtain parameters of rates $k_i$ and diffusion coefficient $d_w$ that satisfy the stability condition and Turing instability using the `FindInstance` function in Wolfram Mathematica,

2. Starting from this parameter set we obtain representative parameter sets by varying parameters $k_i$ and the diffusion coefficient $d_w$ one at the time. We do by sampling uniformly four values in the range that satisfy the stability and Turing instability conditions leaving all the other parameters unchanged. In total for each network, we consider at least 24 parameter sets.

3. We solve characteristic polynomial for this parameters finding the solutions $\lambda_1(q), \lambda_2(q), \lambda_3(q)$ and select the solution $\lambda_i(q)$ with a positive real part.

4. We check for the existence of the limit of $\lambda_i(q)$ to be greater than zero for $q \to \infty$ with the `Limit` function in Wolfram Mathematica.

5. Finally we find the maximum of $\lambda_i(q)$ and check weather is lower than the limit for $q \to \infty$.

networks where the maximum of $\lambda_i(q)$ is lower than the asymptote for all the parameter sets are categorized as noise-amplifying.

**D. Criteria for Turing network classification.** Since we observe that network can respect multiple conditions in table S4 for different parameter sets we use the following criteria for network classification:

| | Cond. Table S4 | | | | |
|---|---|---|---|---|---|
| **Network Type** | **1.** | **2.** | **3.** | **4.** | **Network color** |
| Turing | ✓ | ✗ | ✗ | ✗ | yellow |
| Wave | ✗ | ✓ | ✗ | ✗ | blue |
| Multifunctional | ✓ | ✓ | ✓ | ✗ | green |
| | ✓ | ✓ | ✗ | ✗ | |
| Noise amplifying | ✓ | ✗ | ✗ | ✓ | red |
| | ✗ | ✓ | ✗ | ✓ | |
| | ✓ | ✓ | ✗ | ✓ | |
| | ✓ | ✓ | ✓ | ✓ | |

**Table S5**

Here, it can be observed that multifunctional networks may or may not have a parameter space region that simultaneously satisfies both Turing and Wave conditions (condition 3 in Table S4). On the other hand, we consider that any network could potentially be noise-amplifying if, for all parameters satisfying conditions 1, 2, and 3, the maximum of the positive eingenvalue is the asymptote for $q \to \infty$, which is ensured by condition 4.

**E. Calculation of Network Robustness.** To estimate the robustness of each network $j$, we quantify the volume of the parameter space that satisfies the diffusion-driven conditions derived from the linear stability analysis. This involves integrating all the diffusion-driven instability conditions $f_j$ and is done by fixing one negative feedback rate at $-1$ and one diffusion coefficient at 1, thereby calculating the relative parameter space. The relative parameter space volume is calculated over the ranges: 0.1 to 10 for reaction rates $k_i$ and 0.001 to 100 for the relative diffusion coefficient ratio $d$.

For each network type, the robustness is computed based on the specific conditions the network satisfies. For Turing, Wave, and Noise Amplifying networks, we calculate the volume of parameter space that satisfies conditions 1, 2, and 4 in Table S4, respectively. In the case of multifunctional networks, which can respect both Turing and Wave conditions, the robustness is calculated as the sum of the volumes satisfying the conditions 1 and 2), ensuring that regions satisfying both conditions are only counted once. This is achieved by subtracting the volume of the parameter space that satisfies condition 3 in Table S4, which corresponds to regions where both conditions are simultaneously fulfilled.

The robustness $R(f_j)$ is thus calculated as the integral over the defined parameter space:

$$R(f_j) = \int_{0.1}^{10} \cdots \int_{0.001}^{100} f(k_i, d)\, dk_i\, dd$$

This integration is performed using the `NIntegrate` function in Wolfram Mathematica considering the the boolean volume of the inequalities with the function `Boole`. For the integration we use a `"LocalAdaptive"` based on the `"GaussKronrodRule"`. The `"LocalAdaptive"` method adapts the integration step locally by subdividing the integration domain in regions where the function shows higher error calculated as the difference from a Gauss quadrature and and Kronrod formula (`"GaussKronrodRule"`) to enhance precision within each subdivision, capturing fine details of the diffusio-driven instability volume. This is crucial for reaction-diffusion systems, where as $d \to 1$, the parameter space often forms narrow, localized regions.

After obtaining $R(f)$ for each network, we standardize the robustness values by dividing each $R(f)$ by the hyper-volume of the whole parameter space considered (i.e., the most robust network), $R(f_{\max})$. The normalized atlas robustness $r(f_j)$ for each network $j$ is then given by:

$$r(f_j) = \frac{R(f_j)}{R(f_{\max})}$$

Finally, the logarithmically scaled robustness value is computed using the formula:

$$\log_{10}[r(f_j) \cdot 0.1]$$

The size of a node representing a network $j$ in the atlas is logarithmically proportional to this scaled robustness value. This approach quantifies the likelihood of a given network achieving diffusion-driven instability with randomly assigned parameters and provides a measure of the network's robustness to parameter changes.

**F. Network distance among minimal Turing networks.** We analyzed all the possible 5376 minimal 6-interactions network topologies by selecting those capable of diffusion-driven instability to generate static, oscillatory or noise-amplyfing patterns. This was done by analyzing the sings of the characteristic polynomial coefficients derived from linear stability analysis, as explained in Material and Methods. We obtain a set 376 minimal networks $\Omega_6^{RD}$ capable of diffusion-driven instability, corresponding roughly to $\mathbf{376/5376 \approx 7\%}$ of all possible networks.

For each pair of networks $(a, b)$, we define the topological distance $D^T(a, b)$ between two networks $a$ and $b$, we use both the connectivity matrices $C$ and the topological matrices $T$ associated with the networks. The topological distance is defined as follows:

$$D^T(a,b) = \frac{1}{2}\left(\sum_{i=1}^{3}\sum_{j=1}^{3}\left|t_{ij}^a - t_{ij}^b\right| + \left|c_{ij}^a - c_{ij}^b\right|\right)$$

where:

- $t_{ij}^a$ and $t_{ij}^b$ are the elements of the topological matrices $T^a$ and $T^b$ for networks $a$ and $b$ respectively.
- $c_{ij}^a$ and $c_{ij}^b$ are the elements of the connectivity matrices $C^a$ and $C^b$ for networks $a$ and $b$ respectively.
- $\left|t_{ij}^a - t_{ij}^b\right|$ represents the absolute difference between the topological elements of the two networks.
- $\left|c_{ij}^a - c_{ij}^b\right|$ represents the absolute difference between the connectivity elements of the two networks.

We analyze the distance between all the network pairs $(a, b)$ in $\Omega_6^{RD}$ and find that that the minimum distance between each network pair is at least 2:

$$D^T(a,b) \geq 2 \quad \forall a \quad \forall b \in \Omega_6$$

For example, consider two networks $a$ and $b$:

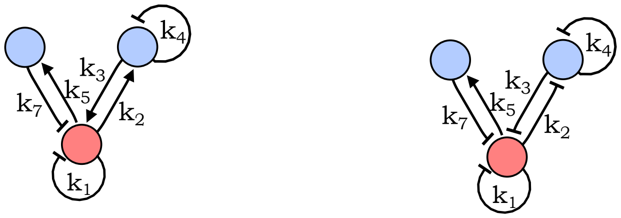

Minimal Turing Networks *a* and *b*

with connectivity matrices $C^a$ and $C^b$ for networks $a$ and $b$:

$$C^a = \begin{pmatrix} 1 & 1 & 1 \\ 1 & 1 & 0 \\ 1 & 0 & 0 \end{pmatrix}, \quad C^b = \begin{pmatrix} 1 & 1 & 1 \\ 1 & 1 & 0 \\ 1 & 0 & 0 \end{pmatrix}$$

and topological matrices $T^a$ and $T^b$ for networks $a$ and $b$ are:

$$T^a = \begin{pmatrix} -1 & 1 & -1 \\ 1 & -1 & 0 \\ 1 & 0 & 0 \end{pmatrix}, \quad T^b = \begin{pmatrix} -1 & -1 & -1 \\ -1 & -1 & 0 \\ 1 & 0 & 0 \end{pmatrix}$$

First, we calculate the element-wise absolute difference between the connectivity matrices:

$$C^{ab} = |C^a - C^b| = \begin{pmatrix} 0 & 0 & 0 \\ 0 & 0 & 0 \\ 0 & 0 & 0 \end{pmatrix}$$

Next, we calculate the element-wise absolute difference between the topological matrices:

$$T^{ab} = |T^a - T^b| = \begin{pmatrix} 0 & 2 & 0 \\ 2 & 0 & 0 \\ 0 & 0 & 0 \end{pmatrix}$$

Then, we sum the differences for both the connectivity and topological matrices:

$$\sum_{i=1}^{3}\sum_{j=1}^{3}\left(|t_{ij}^a - t_{ij}^b| + |c_{ij}^a - c_{ij}^b|\right) = \sum_{i=1}^{3}\sum_{j=1}^{3}(0+2+0+2+0+0+0+0+0) = 4$$

The topological distance $D^T(a,b)$ is calculated as follows:

$$D^T(a,b) = \frac{1}{2}\left(\sum_{i=1}^{3}\sum_{j=1}^{3}\left|t_{ij}^a - t_{ij}^b\right| + \left|c_{ij}^a - c_{ij}^b\right|\right) = \frac{1}{2} \times 4 = 2$$

This means that to transition from a minimal Turing network into another minimal Turing network requires at least two simultaneous reaction rates changes. Since this cannot account for progressive changes and it is an unlikely event in evolution, we decide to include in the atlas extended networks of 7 interactions.

**G. Network distance with extended Turing networks.** To include extended networks, we derive all the possible networks of 7 interactions by setting two rates to zero (showing $R^c$ as an example of the reaction matrix of an extended network $c$):

$$R = \begin{pmatrix} k_1 & k_2 & k_5 \\ k_3 & k_4 & k_6 \\ k_7 & k_8 & k_9 \end{pmatrix} \quad C^c = \begin{pmatrix} 1 & 1 & 1 \\ 1 & 1 & 1 \\ 1 & 0 & 0 \end{pmatrix} \quad R^C = R \circ C^c = \begin{pmatrix} k_1 & k_2 & k_5 \\ k_3 & k_4 & k_6 \\ k_7 & 0 & 0 \end{pmatrix}$$

$$N_3 = \binom{9}{2} = \frac{9!}{2!(9-2)!} = 36$$

As in the previous case the possible topologies for each network are:

$$N_4 = 2^7 = 128$$

Therefore, the total number of networks analyzed in a set of seven rates networks ($\Omega_7$) is:

$$N_3 N_4 = 36 \times 128 = 4608$$

Adding the possible networks of 6 and 7 interactions we obtain:

$$N_1 N_2 + N_3 N_4 = 5376 + 4608 = 9984$$

Of the total of 9984 networks called $\Omega_{67}$ we found that there are 1660 networks that satisfy diffusion-driven instability conditions, called $\Omega_{67}^{RD}$. This corresponds to $\mathbf{1660/9984} \approx \mathbf{16}\%$ of all networks.

We analyze the distance between all the network pairs $(a,b)$ in $\Omega_{67}^{RD}$ and find that that the minimum distance between each network pair is at least 1. This mean that in this extended esamble of networks, it is possible to transition between Turing networks wtih one interaction change at the time.

For example, consider two networks $d$ and $e$:

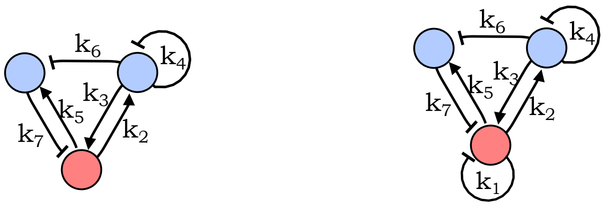

Networks *d* and *e*

With connectivity matrices $C^d$ and $C^e$:

$$C^d = \begin{pmatrix} 0 & 1 & 1 \\ 1 & 1 & 0 \\ 1 & 1 & 0 \end{pmatrix}, \quad C^e = \begin{pmatrix} 1 & 1 & 1 \\ 1 & 1 & 0 \\ 1 & 1 & 0 \end{pmatrix}$$

and topological matrices $T^d$ and $T^e$:

$$T^d = \begin{pmatrix} 0 & 1 & 1 \\ 1 & -1 & 0 \\ -1 & -1 & 0 \end{pmatrix}, \quad T^e = \begin{pmatrix} -1 & 1 & 1 \\ 1 & -1 & 0 \\ -1 & -1 & 0 \end{pmatrix}$$

First, we calculate the element-wise absolute difference between the connectivity matrices:

$$C^{de} = |C^d - C^e| = \begin{pmatrix} 1 & 0 & 0 \\ 0 & 0 & 0 \\ 0 & 0 & 0 \end{pmatrix}$$

Next, we calculate the element-wise absolute difference between the topological matrices:

$$T^{de} = |T^d - T^e| = \begin{pmatrix} 1 & 0 & 0 \\ 0 & 0 & 0 \\ 0 & 0 & 0 \end{pmatrix}$$

Then, we sum the differences for both the connectivity and topological matrices:

$$\sum_{i=1}^{3}\sum_{j=1}^{3}\left(|t_{ij}^d - t_{ij}^e| + |c_{ij}^d - c_{ij}^e|\right) = \sum_{i=1}^{3}\sum_{j=1}^{3}(2+0+0+0+0+0+0+0+0) = 2$$

Finally, the topological distance $D^T(d,e)$ is calculated as follows:

$$D^T(d,e) = \frac{1}{2}\left(\sum_{i=1}^{3}\sum_{j=1}^{3}\left|t_{ij}^d - t_{ij}^e\right| + \left|c_{ij}^d - c_{ij}^e\right|\right) = \frac{1}{2} \times 2 = 1$$

To construct the atlas shown in Figure 1, we build an non directed higher-level graph where each node corresponds to a network in $\Omega_{67}^{RD}$ connected by edges with networks that have topological distance $D^T = 1$. The graph is plotted using the *"Graph"* function of Wolfram Mathematica with a default Spring Electrical Embedding, which treats edges as spring and vertices as charges.

**H. Parameters of simulations in Figure 1F-I.** To perform numerical simulations for the network in the atlas, we devise a simple way to construct a PDE system obtaining parameters that satisfy the analytical diffusion-driven instability conditions, see Material and Methods for a detailed description.

The parameters used for the simulation in Figure 1F-I provided in the following:

**Table S6. Parameters for static Turing network, Figure 1F**

| Parameter | $k_3$ | $k_5$ | $k_6$ | $k_7$ | $k_8$ | $k_9$ | $d_v$ | $d_w$ |
|---|---|---|---|---|---|---|---|---|
| **Value** | $-\frac{1}{4}$ | -1 | 1 | $-\frac{1}{2}$ | -1 | -1 | 1 | 1 |

| Parameter | Value |
|---|---|
| Domain size | 19.19 |
| Total time | 185.83 |
| Time step ($\Delta t$) | $\frac{\text{total time}}{20^7}$ |

Reaction matrix and network diagram:

$$R = \begin{pmatrix} 0 & k_3 & k_7 \\ 0 & 0 & k_8 \\ k_5 & k_6 & k_9 \end{pmatrix}$$

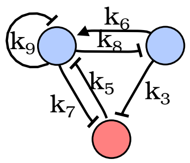

**Table S7. Parameters for Noise amplifying networks, Figure 1G**

| Parameter | $k_1$ | $k_3$ | $k_5$ | $k_6$ | $k_8$ | $k_9$ | $d_v$ | $d_w$ |
|---|---|---|---|---|---|---|---|---|
| **Value** | $\frac{1}{2}$ | $\frac{5}{8}$ | -1 | -1 | 1 | -1 | 1 | 1 |

| Parameter | Value |
|---|---|
| Domain size | 2 |
| Total time | 40 |
| Time step ($\Delta t$) | $\frac{\text{total time}}{20^8}$ |

Reaction matrix and network diagram:

$$R = \begin{pmatrix} k_1 & k_3 & 0 \\ 0 & 0 & k_8 \\ k_5 & k_6 & k_9 \end{pmatrix}$$

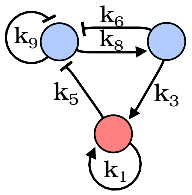

**Table S8. Parameters for Traveling Waves, Figure 1H**

| Parameter | $k_3$ | $k_4$ | $k_5$ | $k_7$ | $k_8$ | $k_9$ | $d_v$ | $d_w$ |
|---|---|---|---|---|---|---|---|---|
| **Value** | $-\frac{67}{128}$ | -1 | -1 | $\frac{17}{32}$ | 1 | $\frac{1}{2}$ | 1 | $\frac{1}{4}$ |

| Parameter | Value |
|---|---|
| Domain size | 36.49 |
| Total time | 217.69 |
| Time step ($\Delta t$) | $\frac{\text{total time}}{20^7}$ |

Reaction matrix and network diagram:

$$R = \begin{pmatrix} 0 & k_3 & 0 \\ 0 & k_4 & k_8 \\ k_5 & k_6 & k_9 \end{pmatrix}$$

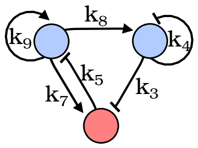

**Table S9. Parameters for Multifunctional network, Figure 1I**

**Static Turing patterns Parameters**

| **Parameter** | $k_3$ | $k_4$ | $k_5$ | $k_6$ | $k_7$ | $k_8$ | $k_9$ | $d_v$ | $d_w$ |
|---|---|---|---|---|---|---|---|---|---|
| **Value** | $\frac{1}{3}$ | $\frac{1}{2}$ | -1 | -1 | -$\frac{1}{9}$ | 1 | -1 | 1 | 1 |

| **Parameter** | **Value** |
|---|---|
| Domain size | 16.25 ç |
| Total time | 852.54 |
| Time step ($\Delta t$) | $\frac{\text{total time}}{20^7}$ |

**Traveling Wave Parameters**

| **Parameter** | $k_3$ | $k_4$ | $k_5$ | $k_6$ | $k_7$ | $k_8$ | $k_9$ | $d_v$ | $d_w$ |
|---|---|---|---|---|---|---|---|---|---|
| **Value** | $\frac{1}{16}$ | 1 | -1 | -2 | $-\frac{5}{64}$ | 1 | $-\frac{3}{2}$ | 1 | 8 |

| **Parameter** | **Value** |
|---|---|
| Domain size | 49.49 |
| Total time | 161.94 |
| Time step ($\Delta t$) | $\frac{\text{total time}}{20^7}$ |

Reaction matrix and network diagram:

$$R = \begin{pmatrix} 0 & k_3 & k_5 \\ 0 & k_4 & k_8 \\ k_7 & k_6 & k_9 \end{pmatrix}$$

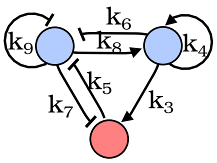

### 2. Diffusion constrains and Phase calculation (Figure 2)

For each network topology in the atlas shown in Figure 1B, as well as for each regulatory logic network in the compressed atlas, we can calculate the type of diffusion constraint. This is possible because, as demonstrated in (8), it is the regulatory logic of the network—defined by the set of network cycle signs—that determines the diffusion constraints of the network (Figure S4).

The calculation of the relative phase, however, can be done only for networks in the atlas in Figure 1B, but not for regulatory logic networks in the compressed atlas shown in Figure 3C. Indeed, the relative phase of the pattern is determined by the specific signs of the regulatory interactions. In the compressed atlas, a set of cycle signs can be implemented by different sets of regulatory interactions. For instance, a positive cycle of size 2 can be implemented either by mutual activation or mutual inhibition, leading to in-phase and out-of-phase patterns, respectively. Therefore, each node in the cycle atlas corresponds to a set of four network topologies, each generating one of the four relative phases between reactants: Phase 1, Phase 2, Phase 3, and Phase 4, shown in Figure 2A.

**A. Classification of diffusion constrain Type.** Each network in the atlas is classified as Type I, Type II, or Type III by evaluating the instability conditions derived from linear stability analysis. First, we simplify the instability conditions by normalizing the values using one of the diffusion coefficients. Specifically, we substitute the value of the diffusion coefficient $d_v$ as $\frac{d_v}{d_v} = 1$ and the value of the diffusion coefficient $d_w$ as $\frac{d_w}{d_v} = d$.

---

**Type III**

If the instability conditions hold true for any value of $d$, the network is classified as a Type III network.
For example, in the network 3 in Figure 2B:

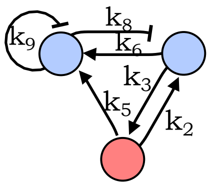

Example of a Type III Network (3 in Figure 3B)

The instability conditions are $d\, k_2\, k_3 > 0$. Given the signs of the reaction rates and that $d > 0$ always, the instability condition simplifies to $True$.

---

### Type II

Next, networks are classified as Type II if the combined stability and instability conditions can be satisfied when $d_v = d_w$ or, using our simplification, if the conditions can be satisfied when $d = 1$.
For example, in the network 2 in Figure 2B:

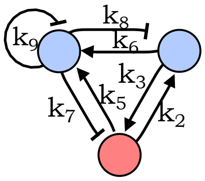

Example of a Type II Network (2 in Figure 3B)

The instability conditions are:

$$dk_2k_3 + k_5k_7 > 0 \vee k_2k_3k_9 < k_2k_6k_7 + k_3k_5k_8$$

. Reducing these conditions along with the stability conditions, we obtain:

$$k_6k_8 < 0 \wedge k_5k_7 < 0 \wedge 0 < k_2k_3 < -k_5k_7 - k_6k_8 \wedge k_2k_6k_7 < 0 \wedge k_3k_5k_8 < 0 \wedge \frac{k_2k_6k_7 + k_3k_5k_8}{k_2k_3} < k_9 < 0 \wedge d > -\frac{k_5k_7}{k_2k_3}$$

These conditions hold even when $d \leq 1$, so the network is classified as a Type II network.

---

### Type I

Otherwise, if the conditions can be only when $d > 1$, the network is classified as Type I.
For example, in the network 1 in Figure 2B:

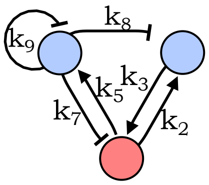

Example of a Type I Network (1 in Figure 3B)

For this network the stability conditions are:

$$\mathrm{k_7k_9 > -k_3k_8 \wedge 0 < k_2k_3 < \frac{k_3k_5k_8}{k_9}}$$

while instability conditions are:

$$\mathrm{d > -\frac{k_5k_7}{k_2k_3}}$$

Together stability and instability conditions can be satisfied when:

$$\mathrm{k_7 < -\frac{k_3k_5k_8}{k_9} \wedge d > -\frac{k_7k_9}{k_3k_8} \wedge -\frac{k_5k_7}{d} < k_2k_3 < \frac{k_3k_5k_8}{k_9}}$$

From which we infer that $\mathrm{d} > 1$ because from the stability conditions we know that $\mathrm{k_7k_9 > -k_3k_8}$.

**B. Relative pattern Phase calculation.** Each network in the atlas generates a relative phase pattern that can be categorized into four distinct configurations:

Phase 1: All three nodes in-phase
Phase 2: Node $v$ out-of-phase, $u$ and $w$ in-phase
Phase 3: Node $u$ out-of-phase, $v$ and $w$ in-phase
Phase 4: Node $w$ out-of-phase, $u$ and $v$ in-phase

To predict the phase pattern generated by a network, we analyze the relative sign of the eigenvectors associated with the eigenvalue that promotes diffusion-driven instability. For each network, we obtain a set of parameters ($k_1..k_9$) with the *FindInstance* command in Wolfram Mathematica that satisfy the diffusion-driven instability conditions. Each parameter $k_i$ is varied within its instability range, and we calculate the eigenvectors $E(\lambda(q_{\max})) = (E_u(q_{\max}), E_v(q_{\max}), E_w(q_{\max}))$ associated with the positive eigenvalue.

For each case, we determine the relative signs of the eigenvectors as follows:

$$E_{u,u} = \frac{E_u(q_{\max})}{E_u(q_{\max})} = 1,\ E_{v,u} = \frac{E_v(q_{\max})}{E_u(q_{\max})},\ E_{w,u} = \frac{E_w(q_{\max})}{E_u(q_{\max})}$$

The relative sign of the phase vector $\varphi = (E_{v,u}, E_{w,u})$ determines the phase of the periodic patterns:

Phase 1, $\varphi_1 : \varphi = (+,+)$
Phase 2, $\varphi_2 : \varphi = (-,+)$
Phase 3, $\varphi_3 : \varphi = (-,-)$
Phase 4, $\varphi_4 : \varphi = (+,-)$

These values of $\varphi$ can be plotted as a function of $k_i$ to identify possible multiphase networks. More details are provided in the Materials and Methods section.

**C. Characterization of multiphase networks .** In the reduced atlas shown in Figure 2A, we observe the presence of multiphase networks where the signs of eigenvectors can change depending on the parameters. We identify these networks through a combination of analytical and numerical approaches, calculating the signs of the eigenvectors for different values of all parameters $k_i$ within the ranges identified by linear stability analysis that give rise to a Turing instability. The proportion of parameters that give rise to a relative pattern phase is calculated with a Multiple integral.

We observed that networks exhibiting multiphase behavior possess two destabilizing feedbacks. Notably, none of these networks are Type I networks.

For example, consider the multiphase network illustrated in Figure 2F:

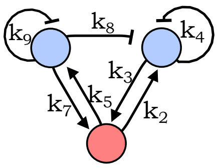

Multiphase network in Figure 2F

The coefficients $a_3(q)$ for this network is:

$$a_3 = k_4k_5k_7 + k_9k_2k_3 - k_3k_5k_8 - d_wq^2\mathbf{k_2\ k_3} - d_vq^2\mathbf{k_5\ k_7}$$

Where $\mathbf{k_2\ k_3}$ and $\mathbf{k_5\ k_7}$ are highlightd in green as the two feedback that can make $a_3(q) < 0$ for a $q > 0$ to promote the formation of a static Turing pattern with different phases.

Code written in Wolfram Mathematica calculates the volume of the parameter space via a numerical multiple integral, with each rate ranging between 0 and 1 if positive, or between -1 and 0 if negative. To simplify, we assign -1 to one of the negative rates and 1 to both diffusions. The integral checks the different possible phases by evaluating the four options:

$$V(\varphi_1) = \int_0^1 ... \int_0^1 \int_{-1}^0 ... \int_{-1}^0 E_{v,u}(k_i,k_j) > 0 \, dk_1...dk_i \, dk_1...dk_j \quad + \int_0^1 ... \int_0^1 \int_{-1}^0 ... \int_{-1}^0 E_{w,u}(k_i,k_j) > 0 \, dk_1...dk_i \, dk_1...dk_j$$
$$V(\varphi_2) = \int_0^1 ... \int_0^1 \int_{-1}^0 ... \int_{-1}^0 E_{v,u}(k_i,k_j) < 0 \, dk_1...dk_i \, dk_1...dk_j \quad + \int_0^1 ... \int_0^1 \int_{-1}^0 ... \int_{-1}^0 E_{w,u}(k_i,k_j) > 0 \, dk_1...dk_i \, dk_1...dk_j$$
$$V(\varphi_3) = \int_0^1 ... \int_0^1 \int_{-1}^0 ... \int_{-1}^0 E_{v,u}(k_i,k_j) < 0 \, dk_1...dk_i \, dk_1...dk_j \quad + \int_0^1 ... \int_0^1 \int_{-1}^0 ... \int_{-1}^0 E_{w,u}(k_i,k_j) < 0 \, dk_1...dk_i \, dk_1...dk_j$$
$$V(\varphi_4) = \int_0^1 ... \int_0^1 \int_{-1}^0 ... \int_{-1}^0 E_{v,u}(k_i,k_j) > 0 \, dk_1...dk_i \, dk_1...dk_j \quad + \int_0^1 ... \int_0^1 \int_{-1}^0 ... \int_{-1}^0 E_{w,u}(k_i,k_j) < 0 \, dk_1...dk_i \, dk_1...dk_j$$

Here, $k_i$ represents the set of positive parameters, and $k_j$ represents the set of negative ones. The integral will be 0 if the evaluated phase is not possible for that network. For typical minimal Turing networks, only one phase per network has an integral that is non-zero. Multiphase extended networks have two phases with an integral different than zero.

To evaluate the parameters contributing to each phase, we normalize the integral values by dividing each by the total value:

$$\Phi = V(\varphi_1) + V(\varphi_2) + V(\varphi_3) + V(\varphi_4), \quad R(\varphi_1) = \frac{V(\varphi_1)}{\Phi}, \; R(\varphi_2) = \frac{V(\varphi_2)}{\Phi}, \; R(\varphi_3) = \frac{V(\varphi_3)}{\Phi}, \; R(\varphi_4) = \frac{V(\varphi_4)}{\Phi}$$

We represent the portion of parameters that give rise to the two phases in multiphase networks by coloring the node with a pie chart, as shown in Figure 2F. Note all Multifunctional nodes are all circular, indicating they are extended networks.

**D. Transition from Phase 1 to Phase 3 (Figure 2D-G).** To test the phase calculated by our theoretical analysis, we performed the simulations showed in Figure 2E-G, focusing on a transition from a network that generate periodic patterns with Phase 1 to a network that generate patterns with Phase 3, passing from a Multiphase network. The simulations were performed with the following parameters:

**Table S10. Parameters for Phase 1 network, Figure 2E**

| Parameter | $k_2$ | $k_3$ | $k_4$ | $k_5$ | $k_8$ | $k_9$ | $d_v$ | $d_w$ |
|---|---|---|---|---|---|---|---|---|
| **Value** | $\frac{1}{2}$ | $\frac{3}{4}$ | -1 | 1 | -1 | -1 | 1 | 1 |

| Parameter | Value |
|---|---|
| Domain size | 14.74 |
| Total time | 320.00 |
| Time step ($\Delta t$) | $\frac{\text{total time}}{10^7}$ |

$$R = \begin{pmatrix} 0 & k_2 & k_5 \\ k_3 & k_4 & 0 \\ 0 & k_8 & k_9 \end{pmatrix}$$

**Table S11. Parameters for Phase 1/3 network, Figure 2F left (Phase 1)**

| Parameter | $k_2$ | $k_3$ | $k_4$ | $k_5$ | $k_7$ | $k_8$ | $k_9$ | $d_v$ | $d_w$ |
|---|---|---|---|---|---|---|---|---|---|
| **Value** | $\frac{1}{4}$ | 1 | -2 | $\frac{3}{4}$ | 0.08 | -3 | -1 | 1 | 1 |

| Parameter | Value |
|---|---|
| Domain size | 5.22 |
| Total time | 3999.82 |
| Time step ($\Delta t$) | $\frac{\text{total time}}{10^7}$ |

$$R = \begin{pmatrix} 0 & k_2 & k_5 \\ k_3 & k_4 & 0 \\ k_7 & k_8 & k_9 \end{pmatrix}$$

**Table S12. Parameters for Phase 1/3 network, Figure 2F right (Phase 3)**

| Parameter | $k_2$ | $k_3$ | $k_4$ | $k_5$ | $k_7$ | $k_8$ | $k_9$ | $d_v$ | $d_w$ |
|---|---|---|---|---|---|---|---|---|---|
| **Value** | $\frac{1}{4}$ | 1 | -2 | $\frac{3}{2}$ | 1 | -3 | -1 | 1 | 1 |

| Parameter | Value |
|---|---|
| Domain size | 15.38 |
| Total time | 149.88 |
| Time step ($\Delta t$) | $\frac{\text{total time}}{10^7}$ |

$$R = \begin{pmatrix} 0 & k_2 & k_5 \\ k_3 & k_4 & 0 \\ k_7 & k_8 & k_9 \end{pmatrix}$$

### 3. Compressed Atlas construction (Figure 3)

In this section, we provide a detailed description of the construction of the compressed Atlas shown in Figure 2. The atlas is constructed by reducing networks in Figure 1B that have the same regulatory logic into a unique node in an atlas of networks represented by cycles. Each node is this compressed atlas is expressed by a set of cycle signs. Cylces are are labeled red if they a negative signs (i.e. negative cycle weight) and green if they have a positive sings (i.e. negative cycle weight). Similar to the normal atlas, each node in labeled (colored) according to its self-organizing behaviour. For each network in the compressed atlas, we derive simpler conditions for diffusion-driven instability by reformulating the coefficients of the characteristic polynomial in terms of cycle, as explained in (5, 8) and in the following.

**Table S13. Parameters for Phase 3 network, Figure 2G**

| Parameter | $k_3$ | $k_4$ | $k_5$ | $k_7$ | $k_8$ | $k_9$ | $d_v$ | $d_w$ |
|---|---|---|---|---|---|---|---|---|
| Value | 2 | -1 | $\frac{1}{2}$ | 1 | -1 | -1 | 1 | 1 |

| Parameter | Value |
|---|---|
| Domain size | 14.66 |
| Total time | 480.00 |
| Time step ($\Delta t$) | $\frac{\text{total time}}{10^7}$ |

$$R = \begin{pmatrix} 0 & 0 & k_5 \\ k_3 & k_4 & 0 \\ k_7 & k_8 & k_9 \end{pmatrix}$$

**A. Compression of the atlas with cycle analysis.** Given a network $i$ in the atlas in Figure 1B, we start by obtaining the adjacency matrix $A^i$ of the associated directed graph. The adjacency matrix is derived by transposing the topological matrix $A^i = (T^i)^\top$.

We then construct the graph associated with each network using the function `AdjacencyGraph` and find all the cycles in the network using the function `FindCycle`. This gives the list of cycles in the graph organized by size, which is a subset of the non-zero cycles among the possible 8 cycles $(c_1, c_2, \ldots, c_8)$ for a three-node network shown in Figure 2A. We find that minimal networks have 4 cycles, while extended networks have 5 cycles.

Next, we calculate the sign of each cycle weight. The weight of a cycle corresponds to the product of the rates $k_i$ that belong to the cycle. The sign of the weight, which we refer to as the sign of the cycle, is obtained by multiplying the elements $a_{ij}$ of the adjacency matrix $A^i$ that are part of the cycle. For each cycle, we obtain 1 if it is a positive cycle, and -1 if it is a negative cycle.

To represent a network in terms of cycle signs (the equivalent of $T^i$ for cycles), we define the regulatory logic for each network $T_c^i = (T_c^{i+}, T_c^{i-})$, where $T_c^{i+}$ is the set of positive cycles in the network and $T_c^{i-}$ is the set of negative cycles.

Given two networks $a$ and $b$, we define the regulatory logic distance of the networks $a$ and $b$ as the size of the difference between $T_c^a$ and $T_c^b$ as:

$$|T_c^a - T_c^b| = |T_c^{a+} - T_c^{b+}| + |T_c^{a-} - T_c^{b-}|$$

We define that two networks have the same regulatory logic if $|T_c^a - T_c^b| = 0$.

For example, consider the following two networks $a$ and $a$ with their topological matrices:

$$T^a = \begin{pmatrix} 0 & 1 & 1 \\ 1 & -1 & 0 \\ -1 & 0 & -1 \end{pmatrix}, \quad T^b = \begin{pmatrix} 0 & -1 & -1 \\ -1 & -1 & 0 \\ 1 & 0 & -1 \end{pmatrix}$$

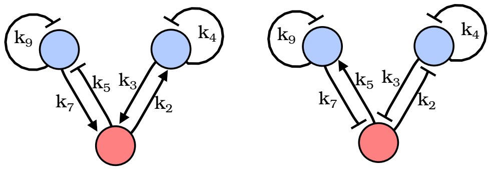

network diagram of networks $a$ and $b$

Performing the cycle analysis for the networks we obtain $T_c^a = \{(c_6), (c_2, c_3, c_5)\}$ and $T_c^b = \{(c_6), (c_2, c_3, c_5)\}$

In both networks, the cycle $c_6$ is positive, either a mutual activation and mutial inhibition and the cycles $c_2, c_3, c_5$ are negative, so they are equivalent in terms of regulatory logic, as shown by $|T_c^a - T_c^b| = 0$.

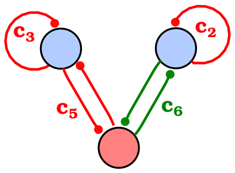

identical regulatory logic for networks $a$ and $b$

By analyzing the regulatory networks of all networks $\Omega_{67}^{RD}$ in the atlas in Figure 1B, which totals 1668, we find that an average of 8 networks reduce to the same regulatory logic, although there are few exceptions. This allows us to compress the atlas in Figure 1B by reducing its size by a factor of 8, resulting in a set $\Omega_{67}^c$ of just **218 networks expressed in terms of cycles**.

In the compressed atlas, where each network $\Omega_{67}^c$ is a node, we can establish direct connections between nodes representing two networks $d$ and $e$ when the size of the difference between their regulatory logic is $|T_c^d - T_c^e| = 1$.

Finally, we calculate for each node the condition for diffusion-driven instability by reformulating the coefficients of the characteristic polynomial $a_1(q)$, $a_2(q)$, and $a_3(q)$ in terms of cycle signs, as explained in (5, 8). Using the same simplified conditions on the signs of the coefficients of the characteristic polynomial presented in Material and Methods and discussed in the next section.

**B. Diffusion-driven instability in terms of cycles.** The cycle analysis presented in (5, 8) allows to rewrite that characteristic polynomial coefficients $a_1(q), a_2(q), a_3(q)$ in terms of network cycle weights and diffusion coefficients. Each characteristic polynomial coefficient can be reformulated as a sum of three terms: a reaction term that depend only on cycles, a reaction+diffusion term that depends on cycles multiplied by diffusion coefficients, and a diffusion term that depend only on diffusion coeffcients, see (5, 8) for details. This is a convenient factorization, because it allows to identify the terms that promote stability within reaction term; and those the promote diffusion-driven instability in the reaction+diffusion term.

An example of the diffusion-driven instability analysis based on cycles is given in the following.

Let's consider a network $a$ represented by the reaction matrix $R^a$ and topology matrix $T^a$:

$$R^a = \begin{pmatrix} 0 & k_2 & k_5 \\ k_3 & k_4 & k_6 \\ k_7 & 0 & 0 \end{pmatrix} \qquad T^a = \begin{pmatrix} 0 & 1 & -1 \\ 1 & -1 & 0 \\ 1 & -1 & 0 \end{pmatrix}$$

and network diagram:

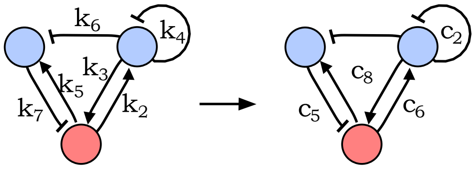

The network contains one cycle $c_2 = k_4$ of length 1, two cycles $c_6 = k_2k_3$ and $c_5 = k_5k_7$ of length 2; and one cycle $c_8 = k_2k_6k_7$ of length 3. The associated regulatory logic is $T^a_c = \{(c_6, c_8), (c_2, c_5)\}$, where $c_6$ and $c_8$ are cycles with a positive weight and the rest have negative weight.

The coefficients of the characteristic polynomial can be rewritten in terms of cyles as:

$$\begin{cases} Coefficients & Reaction & Reaction + Diffusion & Diffusion \\ a_1(q) = & -c_2 & +0 & +d_vq^2 + d_wq^2 \\ a_2(q) = & -c_6 - c_5 & -d_wq^2c_2 & +d_vd_wq^4 \\ a_3(q) = & c_2c_5 - c_8 & -d_wq^2c_6 - d_vq^2c_5 & 0 \end{cases}$$

The stability conditions (i.e. $\triangle_1(0) > 0$ and $\triangle_2(0) > 0$ and $\triangle_3(0) > 0$) can be derived as:

$$\frac{c_8}{c_5} > c_2 > -\frac{c_8}{c_6}$$

The diffusion driven instability conditions for the formation of static Turing patterns (i.e. $a_3(q) < 0$ for $q > 0$) can be derived as:

$$c_5\ d_v + c_6\ d_w > 0$$

Together stability and diffusion-driven instability conditions can be satisfied only when $d_w > d_v$, giving rise to a Type I network. Setting two rates to specific values, for example $c_2 = -1$ and $c_8 = 1$, and considering the ration between diffusion coefficient $d = \frac{d_w}{d_v}$ we can plot the parameter space that give rise to the diffusion driven instability:

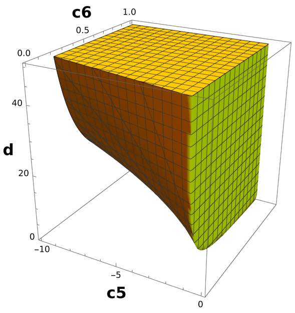

Turing parameter space for network $a$ in terms of cycles $c_5$ and $c_6$, and diffusion ratio $d$

**C. Identification of the cycles that promote diffusion-driven instability.** To identify the network modules (cycles) shown in Figure 3F that produce diffusion-driven instability, we analyze the signs of the different terms in the coefficients of the characteristic polynomial, identifying which cycles contribute negatively to the coefficient. This involves checking the signs of the different terms in the coefficient $a_i(q)$ by assigning -1 to negative cycles and 1 to positive cycles. For each negative term identified, we define the destabilizing cycles as those that contribute to making the term negative.

In the example shown in Section 3B, we analyze the coefficients of the characteristic polynomial expressed in terms of cycles and find that the network can generate static Turing patterns by allowing $a_3(q) < 0$.

The coefficient $a_3(q)$ in terms of cycles is:

$$\begin{array}{cccc} \text{Coefficients} & \text{Reaction} & \text{Reaction+Diffusion} & \text{Diffusion} \\ a_3(q) & = c_2 c_5 - c_8 & -d_w q^2 \mathbf{c_6} - d_v q^2 \mathbf{c_5} & 0 \end{array}$$

The part of the coefficient $a_i(q)$ that can promote diffusion-driven instability is always the *Reaction + Diffusion* part. In this case, there are two terms that can contribute negatively in the *Reaction + Diffusion* part: $-d_w q^2 \mathbf{c_6}$ and $-d_v q^2 \mathbf{c_5}$. Both have negative signs, which means that only the first term with a positive cycle $-d_w q^2 \mathbf{c_6}$ (shown in green) can be overall negative. Therefore, the cycle $c_6$ is the destabilizing cycle.

The Venn diagram in Figure 3F shows which cycles are identified as destabilizing cycles for each type of diffusion-driven instability.

**D. Obtaining a Jacobian from Turing conditions in terms of cycles.** Although the conditions in terms of cycles are equivalent to those derived using an explicit Jacobian and Diffusion matrix, cycle values themselves cannot be directly simulated. As mentioned in the previous section, the regulatory logic of a network expressed as a set of cycle signs $T_c$ corresponds to four different topologies in terms of rates represented by a reaction matrix $R$ and four topology matrices $T^i$.

The correspondence between $T_c$ and $R$ is straightforwardly given by the definition of the cycle weights $c_i$:

$$\begin{array}{ccc} \text{Size 1} & \text{Size 2} & \text{Size 3} \\ c_1 = k_1 & c_4 = k_6\ k_8 & c_7 = k_3\ k_5\ k_8 \\ c_2 = k_4 & c_5 = k_5\ k_7 & c_8 = k_2\ k_6\ k_7 \\ c_3 = k_9 & c_6 = k_2\ k_3 & \end{array}$$

Cycle weight definition

A set of reaction rates to populate the $R$ matrix that satisfies the conditions derived in terms of cycles can be obtained by substituting each cycle with its weight definition and using the *FindInstance* method of Wolfram Mathematica.

The different network topologies $T_i$ associated with the possible sets of reaction rate signs that can give rise to specific cycle signs can be listed according to the following table:

$$\begin{array}{ccc} \text{Size 1} & \text{Size 2} & \text{Size 3} \\ c^+ = (+) & c^+ = (+,+) \text{ or } (-,-) & c^+ = (+,+,+) \text{ or } (-,-,+) \text{ or } (+,-,-) \text{ or } (-,+,-) \\ c^- = (-) & c^- = (+,-) \text{ or } (-,+) & c^- = (-,-,-) \text{ or } (+,+,-) \text{ or } (-,+,+) \text{ or } (+,-,+) \end{array}$$

Possible implementation for positive $c^+$ or negative $c^-$ cycles by set of reaction rate signs

Where it must also be ensured that the sign for a rate $k_{2,3,5,6,7,8}$ can simultaneously satisfy the sign of both cycles of size 2 and size 3 where the rate appears.

### 4. Transitions in the compressed atlas (Figure 4 and S5)

In this section we provide details of the networks analyzed and simulated in the three transition shown in Figure 4 and Figure S5.

**A. Transition from static Turing to Traveling waves (Figure 4B).** In this section, we analyze the path shown in Figure 4B, illustrating a transition from a static Turing network to a traveling wave network through a Multifunctional network. This analysis exploits the cycle analysis discussed in section 3B-C. We detail the coefficients of the characteristic polynomial and observe how the behavior of the network changes with the addition or deletion of specific cycles representing the regulatory feedbacks.

**Static Turing Network:**

$$\begin{cases} Coefficients & Reaction & Reaction + Diffusion & Diffusion \\ a_1(q) & -c_3 & +0 & +d_vq^2 + d_wq^2 \\ a_2(q) & -c_6 - c_5 & -d_wq^2c_3 & +d_vd_wq^4 \\ a_3(q) & c_3c_6 - c_7 & -d_wq^2\mathbf{c_6} - d_vq^2c_5 & +0 \end{cases}$$

Instability conditions: $d_vc_5 + d_wc_6 > 0 \vee c_3c_6 < c_7$
Signs of cycles: $c_3 < 0, c_5 < 0, c_6 > 0, c_7 > 0$
In this case, $a_3 < 0$ due to the cycle $c_6$, and $a_2 > 0$, thus this network exhibits only a static Turing behavior.

**Table S14. Parameters for static Turing network, Figure 4B left**

| Parameter | $k_2$ | $k_3$ | $k_5$ | $k_7$ | $k_8$ | $k_9$ | $d_v$ | $d_w$ |
|---|---|---|---|---|---|---|---|---|
| Value | -1 | -1 | $-\frac{3}{2}$ | 1 | $-\frac{3}{2}$ | -1 | 1 | 2 |

| Parameter | Value |
|---|---|
| Domain size | 16.37 |
| Total time | 634.56 |
| Time step ($\Delta t$) | $\frac{\text{total time}}{10^7}$ |

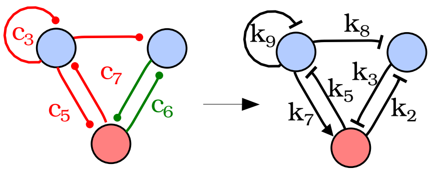

**Traveling Wave Network:**

$$\begin{cases} Coefficients & Reaction & Reaction + Diffusion & Diffusion \\ a_1(q) & -c_2 - c_3 & +0 & +d_vq^2 + d_wq^2 \\ a_2(q) & c_2c_3 - c_5 & -d_wq^2\mathbf{c_2} - d_vq^2c_3 & +d_vd_wq^4 \\ a_3(q) & c_2c_5 - c_7 & -d_vq^2c_5 & +0 \end{cases}$$

Instability conditions: $(c_2c_3 < c_5 \wedge d_wc_2 + d_vc_3) \vee (2d_vd_wc_2c_3 < d_w^2c_2^2 + d_v^2c_3^2 + 4d_vd_wc_5 \wedge d_wc_2 + d_vc_3 \geq 0)$
Signs of cycles: $c_3 < 0, c_5 < 0, c_2 > 0, c_7 > 0$
In this case, $a_2 < 0$ due to the cycle $c_2$ and $a_3 > 0$, thus this network exhibits a traveling wave behavior.

**Table S15. Parameters for Traveling Wave network, Figure 4B right**

| Parameter | $k_3$ | $k_4$ | $k_5$ | $k_7$ | $k_8$ | $k_9$ | $d_v$ | $d_w$ |
|---|---|---|---|---|---|---|---|---|
| Value | $-\frac{11}{8}$ | $\frac{7}{8}$ | -1 | 1 | -1 | -1 | 1 | 4 |

| Parameter | Value |
|---|---|
| Domain size | 251.33 |
| Total time | 62.34 |
| Time step ($\Delta t$) | $\frac{\text{total time}}{10^7}$ |

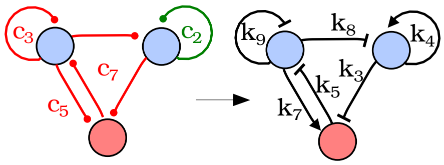

### B. Analysis of the Multifunctional network (Figure 4B-E).

**Multifunctional Network:**

$$\begin{cases} Coefficients & Reaction & Reaction + Diffusion & Diffusion \\ a_1(q) & -c_2 - c_3 & +0 & +d_vq^2 + d_wq^2 \\ a_2(q) & c_2c_3 - c_6 - c_5 & -d_wq^2\mathbf{c_2} - d_vq^2c_3 & +d_vd_wq^4 \\ a_3(q) & c_3c_6 + c_2c_5 - c_7 & -d_wq^2\mathbf{c_6} - d_vq^2c_5 & +0 \end{cases}$$

Instability conditions for Turing: $d_vc_5 + d_wc_6 > 0 \vee c_2c_5 + c_3c_6 < c_7$

Instability conditions for Wave: $(c_2c_3 < c_5 + c_6 \wedge d_wc_2 + d_vc_3) \vee (c_2c_3 < \frac{(d_wc_2+d_vc_3)}{4d_vd_w} + c_5 + c_6 \wedge d_wc_2 + d_vc_3 \geq 0)$
Signs of cycles: $c_3 < 0, c_5 < 0, c_2 > 0, c_6 > 0, c_7 > 0$

For large values of $c_6$, the system exhibits Turing behavior, while for large values of $c_2$, it exhibits traveling wave behavior. We can calculate this numerically how the real (Turing) and complex (Traveling waves) maximum eigenvalue $\lambda_{\max}$ change when $c_6$ or $c_2$ are varied. This is done by finding numerically the maximum of the blue and red eigenvalue curves in Figure 4D, obtaining the following graph:

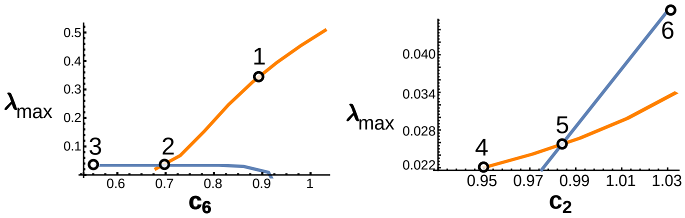

Maximum real part of the real eigenvalue (orange) and complex eigenvalues (blue) as $c_6$ or $c_2$ are varied. Points 1,2,3 correspond to the three graphs in Figure 4D with corresponding simulations in Figure 4E. Points 4,5,6 show the maximum eigenvalue of a similar modulation obtained changing $c_2$ rather than $c_6$.

The graph on the left is obtained by varying $c_6$ with $c_2 = 1$, and the graph on the right is obtained by varying $c_6$ setting $c_6 = 0.65$. The rest of parameters have value $c_3 = -16$, $c_5 = -\frac{69}{4}$, $c_7 = -32$, $d_v = 1$, $d_w = 32$;

**Table S16. Parameters for Multifunctional network, Figure 4E**

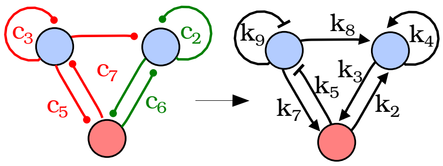

(1 Turing)

| Parameter | $k_2$ | $k_3$ | $k_4$ | $k_5$ | $k_7$ | $k_8$ | $k_9$ | $d_v$ | $d_w$ |
|---|---|---|---|---|---|---|---|---|---|
| **Value** | 1 | 0.9 | 1 | $-\frac{69}{4}$ | 1 | 32 | -16 | 1 | 32 |

| Parameter | Value |
|---|---|
| Domain size | 15.22 |
| Total time | 1359.02 |
| Time step ($\Delta t$) | $\frac{\text{total time}}{10^7}$ |

(2 Turing+Waves)

| Parameter | $k_2$ | $k_3$ | $k_4$ | $k_5$ | $k_7$ | $k_8$ | $k_9$ | $d_v$ | $d_w$ |
|---|---|---|---|---|---|---|---|---|---|
| **Value** | 1 | 0.7 | 1 | $-\frac{69}{4}$ | 1 | 32 | -16 | 1 | 32 |

| Parameter | Value |
|---|---|
| Domain size | 16.18 |
| Total time | 1182.44 |
| Time step ($\Delta t$) | $\frac{\text{total time}}{10^7}$ |

(3 Waves)

| Parameter | $k_2$ | $k_3$ | $k_4$ | $k_5$ | $k_7$ | $k_8$ | $k_9$ | $d_v$ | $d_w$ |
|---|---|---|---|---|---|---|---|---|---|
| **Value** | 1 | 0.55 | 1 | $-\frac{69}{4}$ | 1 | 32 | -16 | 1 | 32 |

| Parameter | Value |
|---|---|
| Domain size | 18.30 |
| Total time | 886.25 |
| Time step ($\Delta t$) | $\frac{\text{total time}}{10^7}$ |

As mentioned above a similar multifunctional network modulation can be achieved by changing the stregth of $c_2$. This can be shown by the following examples:

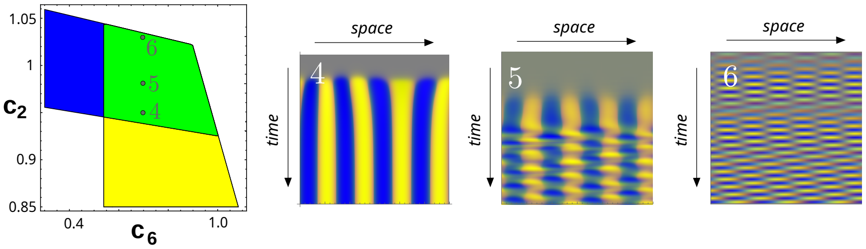

obtained with the following parameters:

(5 Turing+Waves)

| Parameter | $k_2$ | $k_3$ | $k_4$ | $k_5$ | $k_7$ | $k_8$ | $k_9$ | $d_v$ | $d_w$ |
|---|---|---|---|---|---|---|---|---|---|
| **Value** | 1 | 0.65 | 0.95 | $-\frac{69}{4}$ | 1 | 32 | -16 | 1 | 32 |

| Parameter | Value |
|---|---|
| Domain size | 44.77 |
| Total time | 1095.75 |
| Time step ($\Delta t$) | $\frac{\text{total time}}{10^7}$ |

(5 Turing+Waves)

| Parameter | $k_2$ | $k_3$ | $k_4$ | $k_5$ | $k_7$ | $k_8$ | $k_9$ | $d_v$ | $d_w$ |
|---|---|---|---|---|---|---|---|---|---|
| **Value** | 1 | 0.65 | 0.98 | $-\frac{69}{5}$ | 1 | 32 | -16 | 1 | 32 |

| Parameter | Value |
|---|---|
| Domain size | 18.30 |
| Total time | 886.25 |
| Time step ($\Delta t$) | $\frac{\text{total time}}{10^7}$ |

(6 Waves)

| Parameter | $k_2$ | $k_3$ | $k_4$ | $k_5$ | $k_7$ | $k_8$ | $k_9$ | $d_v$ | $d_w$ |
|---|---|---|---|---|---|---|---|---|---|
| **Value** | 1 | 0.65 | 1.03 | $-\frac{69}{4}$ | 1 | 32 | -16 | 1 | 32 |

| Parameter | Value |
|---|---|
| Domain size | 35.63 |
| Total time | 75.88 |
| Time step ($\Delta t$) | $\frac{\text{total time}}{10^7}$ |

**C. Temporal modulation of Multifunctional network, Figure 4F-H.** We performed a 1D and 2D simulation starting from the following parameters that give rise to static Turing pattern within the multifunctional parameter space:

$$c_2 = 1,\ c_3 = -16,\ c_5 = -\frac{69}{4},\ c_6 = 0.85,\ c_7 = -32,\ d_v = 1,\ d_w = 32$$

.

Corresponding to the point $t_0$ in the graph in Figure 4C, which can be implemented by reaction rates:

**Table S17. Parameters for Temporal modulation, Figure 4F-H**

| Parameter | $k_2$ | $k_3$ | $k_4$ | $k_5$ | $k_7$ | $k_8$ | $k_9$ | $d_v$ | $d_w$ |
|---|---|---|---|---|---|---|---|---|---|
| **Value** | 1 | 0.85 | 1 | $-\frac{69}{4}$ | 1 | 32 | -16 | 1 | 32 |

| Parameter | Value |
|---|---|
| Domain size | 34.5 |
| Total time | 600 |
| Time step ($\Delta t$) | $\frac{\text{total time}}{10^7}$ |

$$R = \begin{pmatrix} 0 & k_2 & k_5 \\ k_3 & k_4 & 0 \\ k_7 & k_8 & k_9 \end{pmatrix}$$

And modify the equation of the system to progressively decrease the strength of $c_6$ over time from 0.85 to 0.65 by decreasing $k_3$, from $t_0 = 0$ to $tf = 300$ with $\delta_{k_3} = 6 \cdot 10^{-6}$ as follows:

$$\begin{aligned}
\frac{\partial u}{\partial t} &= k_2\ v + k_5\ w - u^3 \\
\frac{\partial v}{\partial t} &= (k3 - \delta_{k_3}\ \min(\frac{t}{t_f}, 1))\ u + k_4\ v - v^3 + d^v \nabla^2 v \\
\frac{\partial w}{\partial t} &= k7\ u + k_8\ v + k_9\ w - w^3 + d^w \nabla^2 w
\end{aligned}$$

**D. Spatial modulation of Multifuctional network, Figure 4I-M.**

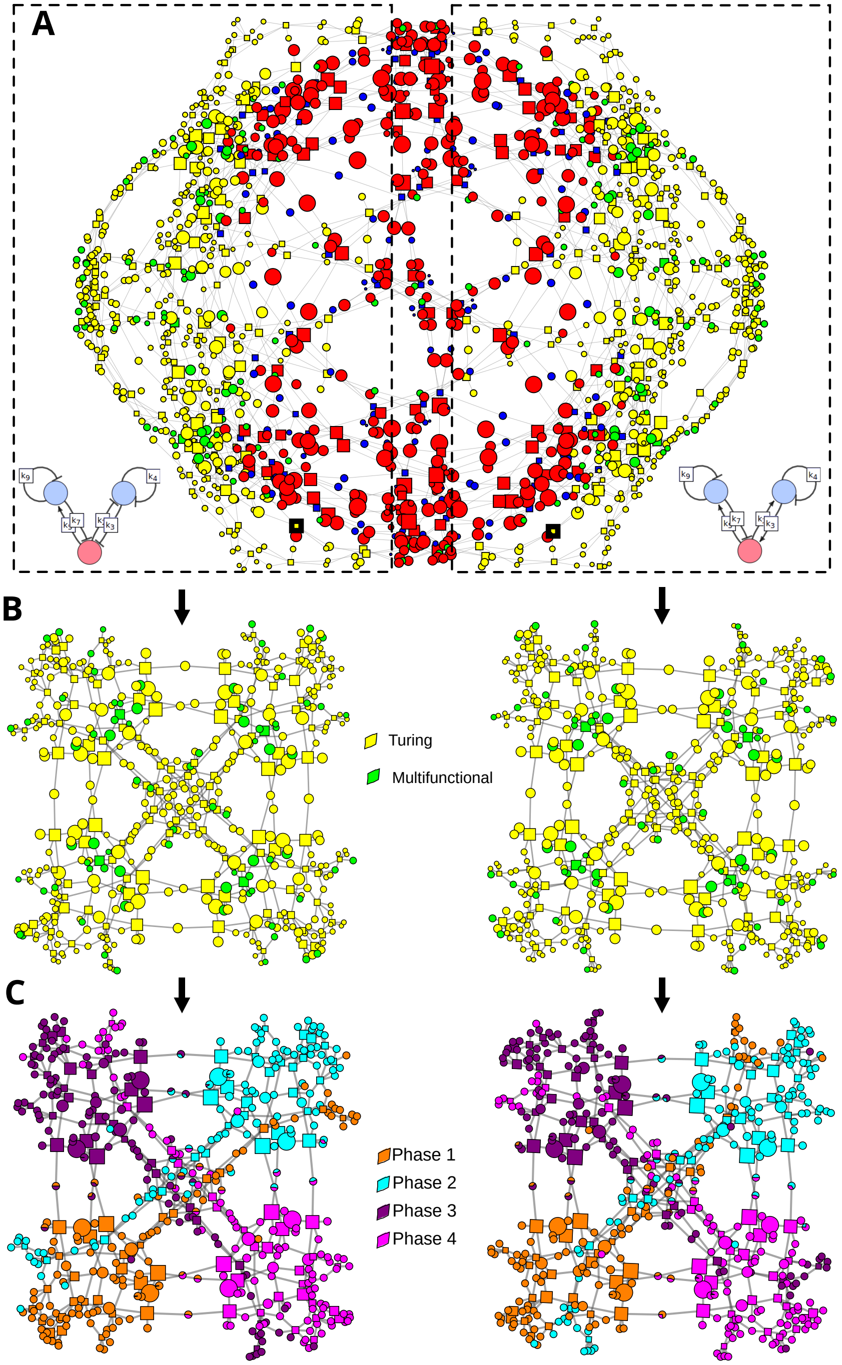

**Fig. S1. Reduced atlas of static Turing networks** The topological atlas shown in Figure 1B exhibits symmetries: similar connectivities and structures from top to bottom and left to right. These symmetries do not represent isomorphic networks, which are excluded in the screening (i.e., networks with the same regulations but an exchange between the diffusible nodes $u$ and $v$). These symmetries represent instead networks where a feedback is exchanged for another feedback with the same overall regulatory signs. For instance, a mutual activation and a mutual inhibition are both a positive regulatory feedback. This example is shown by the two networks at the bottom of panel (A), corresponding to the two nodes highlighted with a thick black border in the atlas. The reduced atlas shown in Figure 3C condenses networks with this type of symmetry into a single network represented in terms of cycle signs. The actual implementation of cycle signs, however, determines the specific relative phase between the periodic patterns formed by the network but does not determine the pattern-forming capabilities of the Turing network. Panel (B) shows that extracting only the subset of networks that can generate static Turing patterns from the atlas in panel (A) results in two separated sub-graphs composed of static Turing networks and multifunctional networks capable of generating static Turing patterns. Our analysis of pattern phases in panel (C) further show that each of these subgraphs is organized into four clusters of networks with a different phase. From a pattern-forming capability perspective, these four clusters are identical, as seen in panel (B). The compressed atlas in Figure 3C condenses networks with these symmetries into the same network represented by a set of cycle signs, as illustrated in Figure 3B.

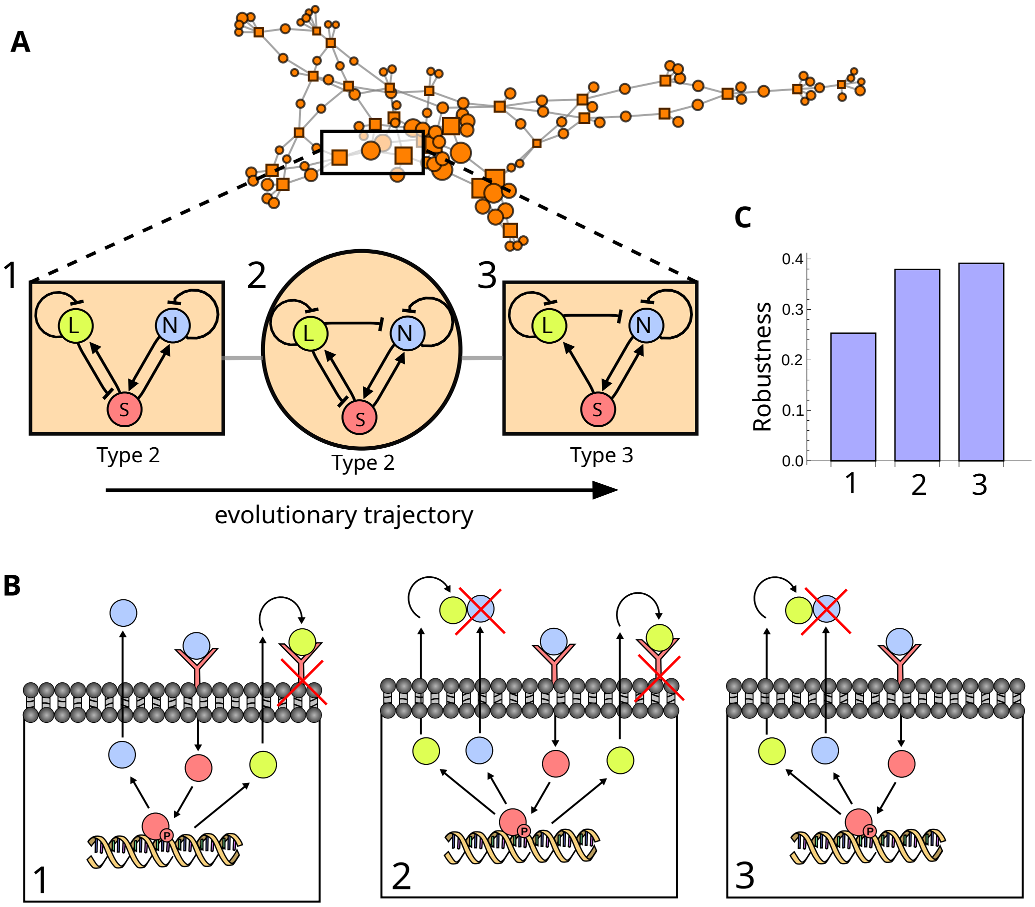

**Fig. S2. Evolutionary trajectory of the Nodal-Lefty network** A) A subset of the static Turing networks in the atlas shown in Figure 3C that generate periodic patterns in-phase for all three reactants (Phase 1). The regulatory topology of the Nodal-Lefty system, which has been proposed to act as a Turing system (10) producing Nodal and Lefty co-expressed in the same place (in-phase), is highlighted by three networks (1, 2, 3) in the subatlas. B-C) The three networks (1, 2, 3) represent different implementations of the inhibition of Nodal by Lefty in the networks. In agreement with our previous analysis in (5), where we only analyzed minimal networks (6 regulatory interactions, square nodes), inhibition of Nodal signaling by interfering with the receptor (network 1) gives rise to a Type II network, while inhibition of Nodal by dimerization with Nodal (network 3) gives rise to a more robust Type III network. In vivo experiments suggest that Lefty inhibits Nodal by binding to its receptor as in network 1 (10), but in vitro evidence also shows that Lefty can bind directly to the Nodal ligand to interfere with its activity (11). In the subatlas, we also find an extended network (network 2 with 7 interactions) that acts as an intermediate between network 1 and 3 and contains both types of inhibitions. The presence of both inhibitions is enough to boost robustness to comparable levels as in network 3. In addition to parameter robustness, network 2 is also robust to loss of one interaction from Lefty as it possess regulatory redundancy .

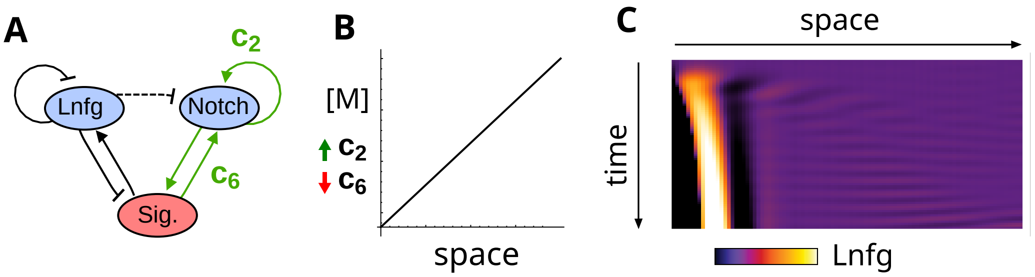

**Fig. S3. Spatial modulation of self-organization in Gastruloids** Spatial modulations of Multifunctional networks, as presented in Figure 4B and I-M, can be used to study multicellular systems where complex patterning dynamics arise from different patterning regimes within the same tissue. In a recent study (12), it was shown that when embedded in Matrigel, Gastruloids exhibit patterning as indicated by the reporter (LfngT2AVenus) with a static peak of Lunatic fringe (Lnfg) expression on one side of Gastruloids and traveling waves on the other. A) The multifunctional network shown in Figure 4B can represent different feedbacks on Notch signaling mediated by Lfng, which is expressed in presomitic mesoderm cells that exhibit Notch signaling oscillations. This inhibition can be cell-autonomous by inhibiting the Notch ligand Delta (13), corresponding to the negative feedback on the non-diffusible node (Sig.), or could be paracrine since Lfng can be secreted extracellularly (14), corresponding to the negative feedback acting on Notch. Similarly, Notch signaling could promote a positive cell-autonomous feedback in response to autocrine Notch signaling (15) ($c_6$), or trigger a positive Notch signaling feedback on neighboring cells via the diffusible Jagged ligand (16) ($c_2$). These interactions have been observed in diverse systems and are speculative within the context of axis formation and somitogenesis, however, they illustrate how the interactions between diffusible and non-diffusible nodes in the atlas can be related to the literature. B-C) A reduction in the strength of autonomous Notch signaling feedbacks $c_6$ and an increase in the strength of non-cell autonomous feedback $c_2$ by a morphogen gradient M, as done in Figure 4I-M, can lead Notch oscillateation closed to the morphgoen source (traveling wave regime) and one static Turing pattern further away from the morphogen. A space-time plot of a simulation shows Lfng expression dynamics that resemble the space-time plot of Lfng quantified in Gastruloids in Figure 2c in (12).

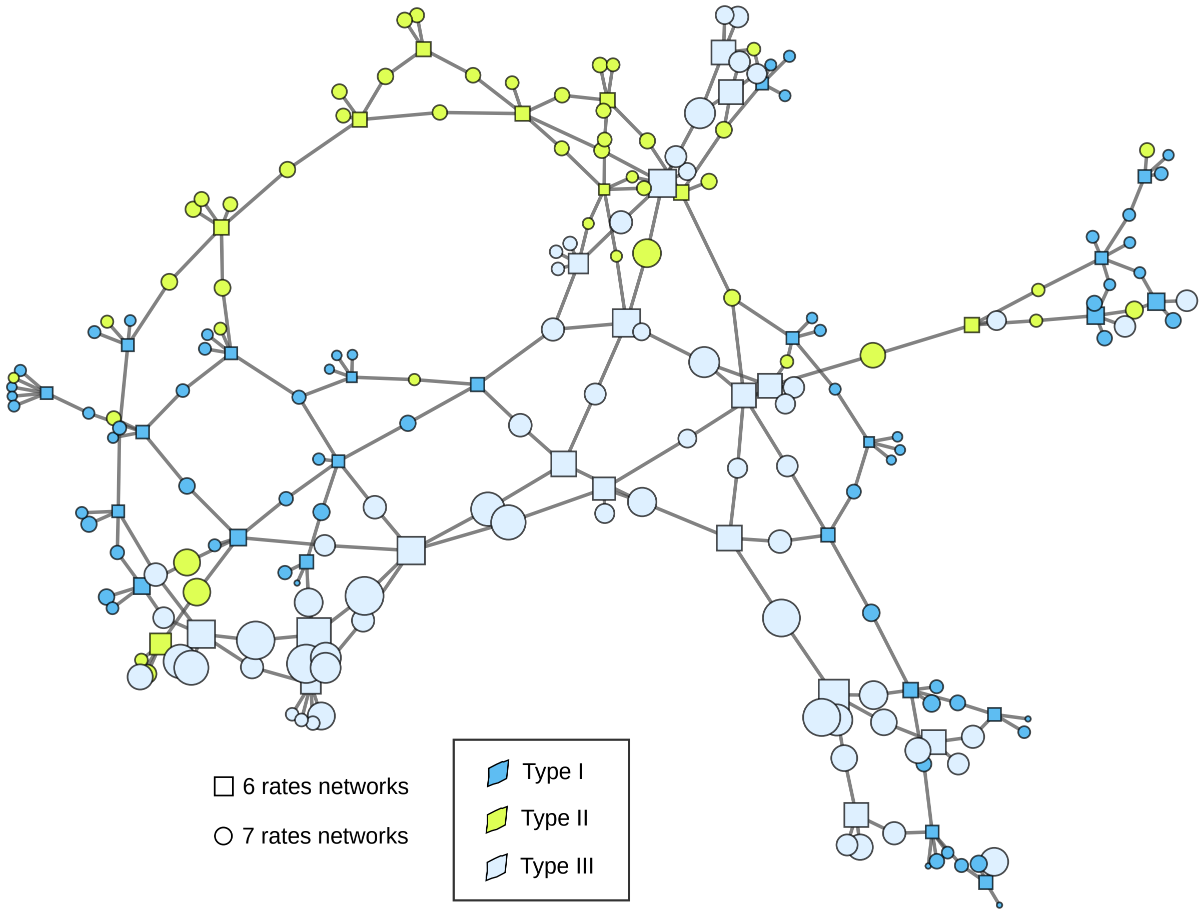

**Fig. S4. Diffusion Constraints in the Compressed Atlas** The compressed atlas in Figure 3C demonstrates that the signs of network cycles are predictive of Turing self-organizing behavior, as evidenced by the clustering of networks with similar self-organizing behaviors in the same region of the compressed regulatory logic graph. Additionally, the atlas confirms that network cycles are predictive of diffusion constraints, as discussed in (8), indicated by the clustering of Type I, II, and III networks within the same topological space in the atlas.

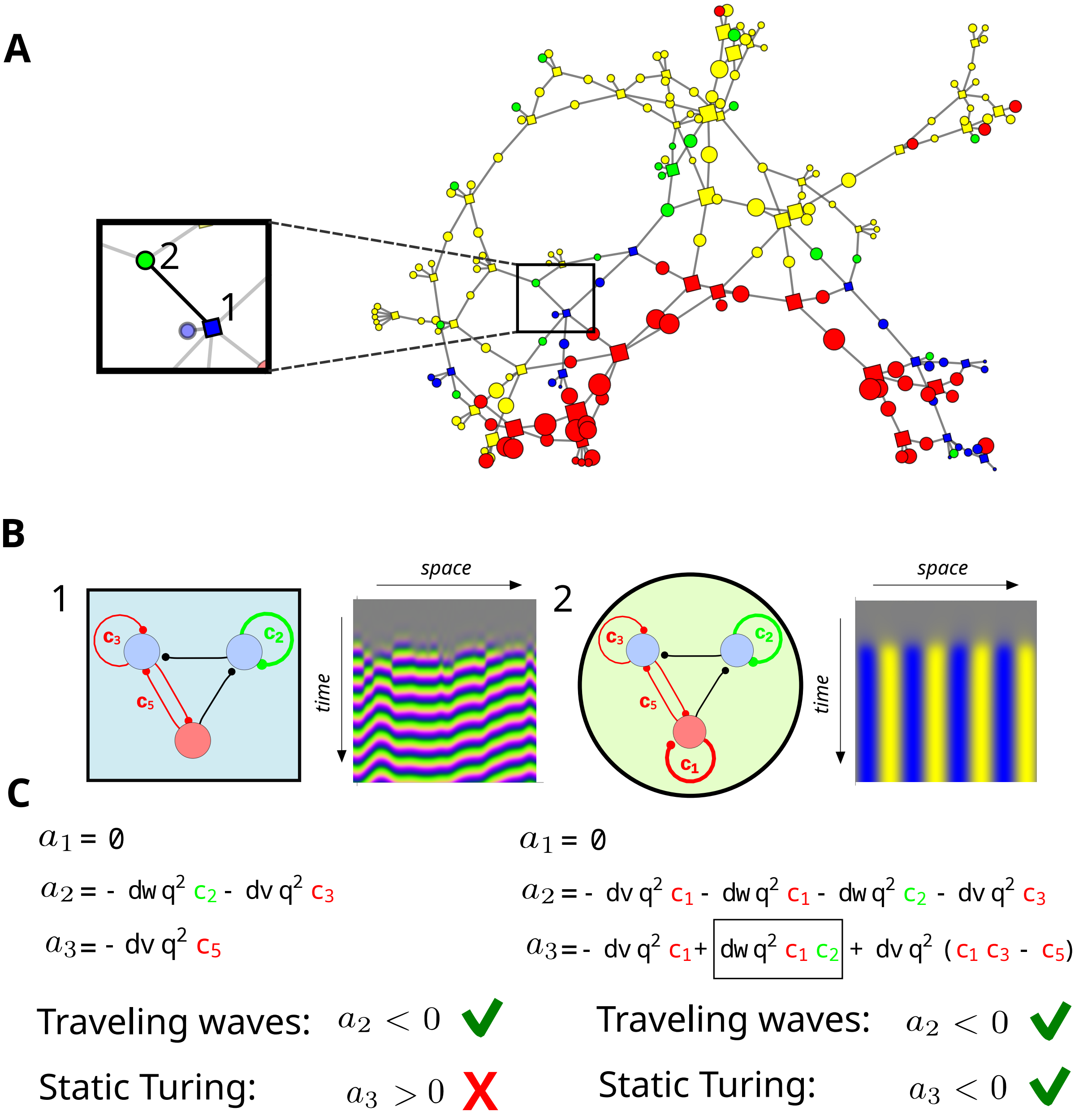

**Fig. S5. Alternative Transition between Traveling Waves and Multifunctional Networks** Many transitions from Traveling Waves networks to Multifunctional and Turing networks in the compressed atlas (Figure 3C) involve the addition of a positive feedback between diffusible and non-diffusible nodes ($c_5$ or $c_6$) along with the auto-regulatory feedback on a diffusible node that promotes traveling waves. This is also evident in the transition shown in Figure 4B. The feedback cycle modularity described in Figure 3F explains this phenomena, since a positive cycle of length two between diffusible and non-diffusible nodes can introduce negative terms in the characteristic polynomial coefficient $a_3$ to make it negative and thus promote static Turing patterns. However, Figure 3F also reveals unexpected cases where static Turing patterns are promoted by a negative self-regulatory feedback on the non-diffusible node ($c_1 < 0$). This is exemplified by the transition highlighted in panels (A) and (B). Panel C illustrates the conditions derived using our symbolic algebraic approach, elucidating this transition. The Traveling Wave network on the left lacks $c_1$ and only has a positive feedback on a diffusible node ($c_2$), which introduces a negative term in $a_2$ and can make it negative. However, there are no negative terms in $a_3$ to make it negative. The addition of the negative feedback on the immobile node ($c_1 < 0$) introduces new terms in $a_2$ and $a_3$, including a negative term coupling $c_1$ and $c_2$, which can make $a_3$ negative and transform the network into a Multifunctional one capable of generating static Turing patterns.
